# Multiplexed in vivo imaging with fluorescence lifetime modulating tags

**DOI:** 10.1101/2024.04.12.589181

**Authors:** Lina El Hajji, France Lam, Maria Avtodeeva, Hela Benaissa, Christine Rampon, Michel Volovitch, Sophie Vriz, Arnaud Gautier

## Abstract

Fluorescence lifetime imaging opens new dimensions for highly multiplexed imaging in live cells and organisms using differences in fluorescence lifetime to distinguish spectrally identical fluorescent probes. Here, we describe a set of fluorescence-activating and absorption-shifting tags (FASTs) capable of modulating the fluorescence lifetime of embedded fluorogenic 4-hydroxybenzylidene rhodanine (HBR) derivatives. We show that changes in the FAST protein sequence can vary the local environment of the chromophore and lead to significant changes in fluorescence lifetime. These fluorescence lifetime modulating tags enabled multiplexed imaging of up to three targets in one spectral channel using a single HBR derivative in live cells and live zebrafish embryo. The combination of fluorescence lifetime multiplexing with spectral multiplexing allowed us to successfully image six targets in live cells, opening great prospects for multicolor fluorescence lifetime multiplexing.

## INTRODUCTION

Fluorescence lifetime microscopy (FLIM) has recently gathered a broad attention in biological imaging thanks to the development of easy-to-use acquisition setups and approaches for data analysis^1^. FLIM leverages the fluorescence lifetime, i.e. the average duration of the excited state, of a fluorescent species for probing a given biological sample. Characterization of the lifetime properties of commonly used fluorescent reporters allowed the use of FLIM in a variety of biological contexts. Combined with Förster Energy Transfer (FRET), FLIM enables detection of protein-protein interactions through changes in the donor fluorescence lifetime^2^. Engineering of fluorescent biosensors with FLIM responsiveness allowed for quantitative imaging of biomolecules of interest through measurements of fluorescence lifetime instead of signal intensity^3^.

One interesting use of FLIM is the possibility to simultaneously visualize several cellular targets relying on their fluorescence lifetime. Multiplexed observation is usually achieved by multicolor fluorescence microscopy using fluorescent tags displaying different spectral properties (different “colors”). The wide excitation and emission spectra of most fluorophores limit however the number of species that can be imaged simultaneously in fluorescence microscopy to four or five. Recently, FLIM based on time-correlated single photon counting (TCSPC) has gathered attention as it opens additional discrimination dimension to distinguish fluorophores with similar colors^4^. Visualization of up to nine fluorescent proteins have been recently achieved using multicolor FLIM^5^, demonstrating the potential of FLIM for highly multiplexed imaging using readily available fluorescent reporters. To expand the possibilities offered by fluorescence lifetime multiplexing, dedicated engineering strategies have been developed using lifetime as a criterium to identify fluorescent proteins^3,6^ and HaloTag-based chemogenetic fluorescent reporters^7^ that display a wider range of fluorescence lifetime differences.

Here, we describe the identification of variants of the fluorescence activating and absorption-shifting tag (FAST) that are able of modulating the fluorescence lifetime of embedded 4-hydroxybenzylidene rhodanine (HBR) derivatives for fluorescence lifetime multiplexing in live cells and organisms. Small proteins of only 14 kDa, FAST and its derivatives are attractive alternatives to fluorescent proteins and self-labeling tags for monitoring gene expression and protein localization with minimal perturbation^8,9^. HBR derivatives only fluoresce when embedded within FAST, enabling wash-free imaging with very high contrast in cells and in multicellular organisms. FASTs have been shown to excel for imaging proteins in anaerobic organisms^10^, and to allow imaging of highly dynamic processes thanks to the extremely rapid formation of the fluorescent assembly^11^. Here, we show that screening of a collection of FAST variants allowed us to identify three variants that can modulate the local chromophore environment, leading to significant change in the fluorescence lifetime of the embedded chromophore. These fluorescence lifetime modulating tags allow the imaging of three targets in one spectral channel using a single fluorogenic chromophore in live mammalian cells and in zebrafish larvae, opening great prospect for highly multiplexed imaging.

## RESULTS

### Identification and characterization of FAST variants for FLIM multiplexing

4-hydroxybenzylidene rhodanine derivatives are push-pull chromophores with fluorescence properties highly dependent on the environment. When free, they are almost non-fluorescent, as they dissipate light energy through ultrafast non-radiative de-excitation pathways. Rotations around the methylene bridge bonds have been proposed to promote internal conversion by conical intersection. When embedded within FAST, their anionic state is stabilized and locked in a planar conformation, leading to fluorescence. Different FAST variants adapted to imaging in different biological contexts have been developed over the last few years, mainly through rational design^12^ and directed evolution^9,13^. Recently, the use of proteins with homology relationship enabled the engineering of new variants, with sequence similarity between 70 to 78 % relative to prototypical FAST, all binding prototypical fluorogens with good affinity and yielding fluorescent assemblies with good brightness in vitro and in cells^14^. All together, these different engineering efforts provide a collection of proteins with a wide diversity of sequences, leading to different photophysical behaviors, and thus hypothetically to different fluorescence lifetimes.

To identify FAST:fluorogen assemblies suitable for fluorescence lifetime multiplexing (**Fig. 1a**), we screened fluorogens able to form tight and bright fluorescent assemblies with the largest possible number of variants. We tested the fluorogens HMBR, HBR-3,5DM and HBR-2,5DM, which form tight (*K*_D_ 10-900 nM) and bright (FQY 20-50 %) green-yellow fluorescent assemblies with the original FAST^8^ (engineered from the photoactive yellow protein (PYP) from *Halorhodospira halophila*), its variants, iFAST^12^, pFAST^9^, oFAST^9^, tFAST^9^ and greenFAST^13^, as well as the six recently-described FAST systems, HboL-FAST, HspG-FAST, RspA-FAST, Ilo-FAST, TsiA-FAST and Rsa-FAST^14^, engineered from six homologs of PYP from *Halomonas boliviensis* LC1 (HboL), *Halomonas* sp. GFAJ-1 (HspG), *Rheinheimera* sp. A13L (RspA), *Idiomarina loihiensis* (Ilo), *Thiorhodospira sibirica* ATCC 700588 (TsiA), and *Rhodothalassium salexigens* (Rsa) (**Supplementary Text 1** and **Supplementary** Figure 1). We imaged each variant:fluorogen assembly in cultured mammalian cells by TCSPC-FLIM (**Supplementary** Figure 2). For the screening, we fitted the fluorescence decays of each assembly with a bi-exponential model and computed the intensity-weighted average fluorescence lifetime (T_i_) (**Supplementary** Figure 2a-c). The use of T_i_ enables to compare systems regardless of their mono or bi-exponential decay behavior. In agreement with their relative fluorescence quantum yields, for a given variant, the T_i_ of the fluorescent assembly gradually increased going from HMBR to HBR-2,5DM and HBR-3,5DM. Interestingly, with the same fluorogen, although displaying very similar fluorescence quantum yields, some FAST variants yielded fluorescent assemblies with clearly distinguishable lifetime signatures. This behavior was already reported for greenFAST and iFAST, which form assemblies with HMBR having similar fluorescence quantum yields (23 % and 22 % respectively), but distinguishable T_i_ values (respectively T_i_ = 1.25 ± 0.01 ns and 1.73 ± 0.01 ns)^13^. Noteworthily, regardless of the fluorogen used, greenFAST was always the variant yielding the lowest T_i_, while TsiA-FAST always gave the highest T_i_ (**Supplementary** Figure 2d). Computing of pairwise lifetime differences enabled the identification of variants suitable for fluorescence lifetime multiplexing. Regardless of the fluorogen, greenFAST displays lifetime difference large enough with most FAST variants for multiplexing by FLIM (ϕλT_i_ > 0.4 ns), the best discrimination being observed with TsiA-FAST. greenFAST and TsiA-FAST displayed a ϕλT_i_ of 0.78 ns with HMBR, 0.89 ns with HBR-2,5DM, and 1.17 ns with HBR-3,5DM. We identified a third variant, oFAST, that forms fluorescent assemblies with fluorescence lifetimes in between those obtained with greenFAST and TsiA-FAST (**Supplementary** Figure 2), opening exciting prospects for fluorescence lifetime multiplexing of three FAST variants in the same spectral channel. In this screening, HBR-2,5DM appeared as the optimal fluorogen for lifetime multiplexing of the three FAST variants, while HBR-3,5DM offered the best lifetime differences for the separation of two FAST variants. Hereafter, greenFAST, oFAST and Tsi-FAST are renamed shortT-FAST, midT-FAST and longT-FAST, respectively, in which the prefix indicates whether they give short, intermediate or long fluorescence lifetimes. Moreover, when designating their assemblies with HBR-2,5DM (resp. HBR-3,5DM), which emits at ∼ 550 nm (resp. 560 nm) within FAST variants, we will use the names shortT550, midT550 and longT550 (resp. shortT560, midT560 and longT560).

**Fig. 1.**
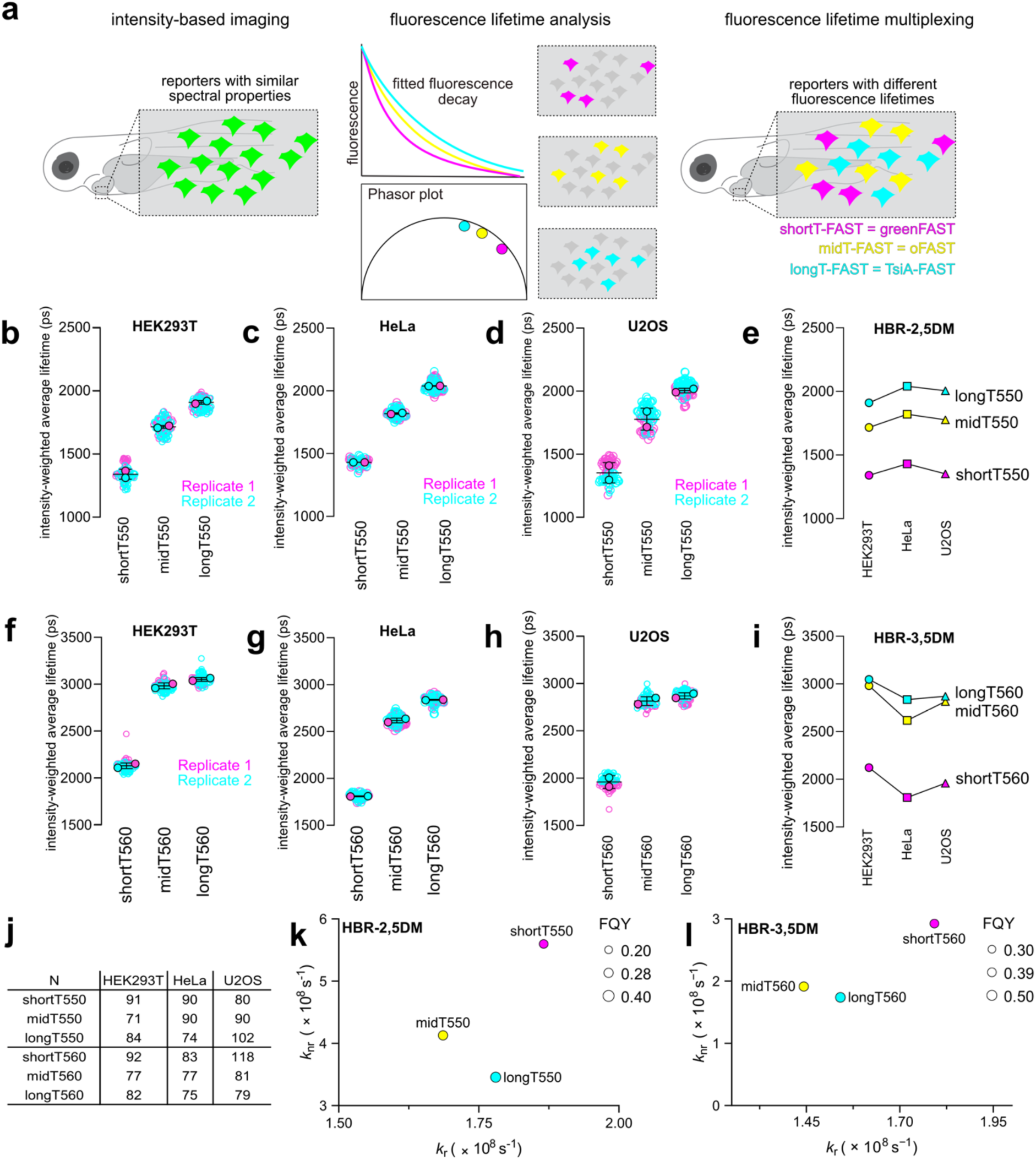
Fluorescence lifetime modulating tags. (**a**) Principle of fluorescence lifetime multiplexing of FAST variants. FAST:fluorogen assemblies with similar spectral properties but different lifetime signatures can be distinguished analyzing their lifetimes. (**b-e**) Intensity-weighted average lifetime distributions of shortT550, midT550 and longT550 in HEK293T cells (**b**), HeLa cells (**c**) and U2OS cells (**d**). N cells from two biological replicates were analyzed (the number N of analyzed cells is given in (**j**)). Each cell is color-coded according to the biological replicate it came from. The solid circles correspond to the mean of each biological replicate. The black line represents the mean ± SD of the two biological replicates. (**e**) Means of intensity-weighted average fluorescence lifetimes of shortT550, midT550 and longT550 in the three cell lines. (**f-i**) Intensity-weighted average lifetime distributions of shortT560, midT560 and longT560 in HEK293T cells (**f**), HeLa cells (**g**) and U2OS cells (**h**). N cells from two biological replicates were analyzed (the number N of analyzed cells is given in (**j**)). Each cell is color-coded according to the biological replicate it came from. The solid circles correspond to the mean of each biological replicate. The black line represents the mean ± SD of the two biological replicates. (**i**) Means of intensity-weighted average fluorescence lifetimes of shortT560, midT560 and longT560 in the three cell lines. (**j**) Number of cells for each set of experiments. (**k,l**) Photophysical properties of FAST variants with HBR-2,5DM (**k**) and HBR-3,5DM (**l**). Each dot corresponds to one FAST:fluorogen assembly according to their radiative and non-radiative decay constant values, and is scaled to the FQY of the assembly.

We next performed an extensive characterization of shortT-FAST, midT-FAST and longT-FAST with HBR-2,5DM and HBR-3,5DM in three different mammalian cell lines, HEK293T, HeLa and U2OS cells (**Fig. 1b-l** and **Supplementary** Figures 3-7). The three variants were expressed as fusions to the histone H2B. We fitted the fluorescence decays with either a monoexponential or a biexponential model, and determined the best fit model using a reduced χ^2^ goodness-of-fit test (**Supplementary** Figures 6 and 7). The variants shortT550 (a.k.a shortT-FAST:HBR-2,5DM) and shortT560 (a.k.a. short-FAST:HBR-3,5DM) clearly displayed biexponential fluorescence decay. The variants midT550 (a.k.a. midT-FAST:HBR-2,5DM) and longT550 (a.k.a. longT-FAST:HBR-2,5DM) displayed rather biexponential decay. For midT560 (a.k.a. midT-FAST:HBR-3,5DM) and longT560 (a.k.a. longT-FAST:HBR-3,5DM), the goodness-of-fit with a monoexponential fit model and a biexponential fit model were very similar. To analyze all the systems the same way and facilitate comparison, we use hereafter T_i_ computed from the results of the biexponential fit. For a given variant, very similar T_i_ values were obtained in the three tested cell lines (**Fig. 1b-d, f-h, j,f**), suggesting that this parameter is robust in various cellular contexts. This study confirmed that, for a given fluorogen, the assemblies with shortT-FAST, midT-FAST and longT-FAST display distinguishable fluorescence lifetimes. With HBR-2,5DM, the three variants show clearly different lifetime signatures in the three cell lines. With HBR-3,5DM, the lifetime difference of the pairs shortT-FAST/midT-FAST and shortT-FAST/longT-FAST was larger than with HBR-2,5DM in all cell lines. However the pair midT-FAST/longT-FAST displays a lifetime difference smaller than with HBR-2,5DM in HEK293T and U2OS cells. In HeLa cells, shortT-FAST and midT-FAST display a difference of lifetime ΔT_i_ = 0.39 ns with HBR-2,5DM and ΔT_i_ = 0.80 ns with HBR-3,5DM. midT-FAST and longT-FAST display a difference of lifetime ΔT_i_ = 0.22 ns with HBR-2,5DM, and ΔT_i_ = 0.22 ns with HBR-3,5DM. Finally, shortT-FAST and longT-FAST display a difference of lifetime ΔT_i_ = 0.61 ns with HBR-2,5DM, and ΔT_i_ = 1.0 ns with HBR-3,5DM. For each variant, the T_i_ values measured in the three different cell lines were close (**Fig. 1e,i**), demonstrating the possibility to distinguish them regardless of the cell type used. We assume that the ability of the three variants to modulate differently the fluorescence lifetime of the embedded fluorogen is related to their different amino acid sequences that modulate the local environment of the chromophore, affecting thus the fate of the excited state. shortT-FAST and midT-FAST share 94 % sequence identity, shortT-FAST and longT-FAST 71 %, and midT-FAST and longT-FAST 74 %. Estimation of the kinetic constants associated to radiative and non-radiative de-excitation suggested that the differences in fluorescence lifetime we observed are mainly due to differences in the non-radiative kinetic constants (**Fig. 1k,l**). While longT-FAST and midT-FAST display close radiative and non-radiative constants regardless of the fluorogen, short-FAST is characterized by a higher non-radiative kinetic constant compared to the two other reporters.

### Fluorescence lifetime multiplexing in mammalian cells

To demonstrate the possibility of performing fluorescence lifetime multiplexing, we first tested pairwise combinations of shortT-FAST, midT-FAST and longT-FAST labeled with either HBR-2,5DM (**Fig. 2**) or HBR-3,5DM (**Fig. 3**). To test each pair, we co-expressed the two variants in HeLa cells in two different cellular localizations, the nucleus and mitochondria, in order to assign one localization to one variant. While the lifetime signature of an emissive species does not depend on its concentration, attention must be given to fluorescence levels to achieve good photon counting, in particular when multiplexing reporters with different cellular brightness^7^. We thus optimized, for each pair, the expression of the two variants to achieve similar fluorescence signals for better photon counting. Various techniques can be used to analyze the FLIM data and separate the contributions of two fluorophores. For each pair of reporters, we compared the photon average arrival time image with the fluorescence decay fitting method and the fit-free phasor approach for separation of the individual variants. Each separation technique bears different advantages and drawbacks: while the fitting approach is widely used to report values of fluorescence lifetime and characterize a given fluorescent reporter, the phasor approach does not require any assumption on the mono, bi– or tri-exponential decay behavior of the reporter’s lifetime and gives a visual representation of the different fluorescent populations present in a given sample. In our case, while the different fluorescent assemblies were undistinguishable in the intensity-only image, we achieved proper discrimination of each variant of a given pair no matter the analysis method (**Fig. 2** and **Fig. 3**), in agreement with their differences in lifetime. For each pair of variants, mitochondria and nucleus displayed different photon average arrival times, in agreement with the presence of two different variants in these two locations (**Fig. 2d,l,t** and **Fig. 3d,l,t**). The phasor representation was used to visualize these two populations, which, for each pair, appear as separate clusters (**Fig. 2e,m,u** and **Fig 3e,m,u**). Segmentation based on the position of these clusters allowed to separate the fluorescence lifetime modulating tags expressed in the mitochondria and the nucleus (**Fig. 2f,n,v** and **Fig. 3f,n,v**). Separate lifetime signatures were also observed on the intensity-weighted average lifetime image obtained through fitting of the fluorescence decays with a biexponential model (**Fig. 2h,p,x** and **Fig3h,p,x**). For all assemblies, the intensity-weighted average lifetime distribution shows two separate bands, allowing the definition of two temporal windows for lifetime-based separation of the two variants (**Fig. 2i,q,y** and **Fig 3i,q,y**). The average lifetime difference remained close to the lifetime difference measured when the variants were expressed alone as H2B fusions in HeLa cells (**Supplementary** Figure 8). Note that, with HBR-3,5DM, the discrimination of midT560 and longT560 was more challenging because of the larger overlap of the two individual lifetime distributions in this context. However, as observed during our initial characterization, shortT560 and longT560 displayed the largest lifetime difference (ΔT_i_ > 1 ns), which makes this pair the best for FLIM multiplexing of two targets.

**Fig. 2.**
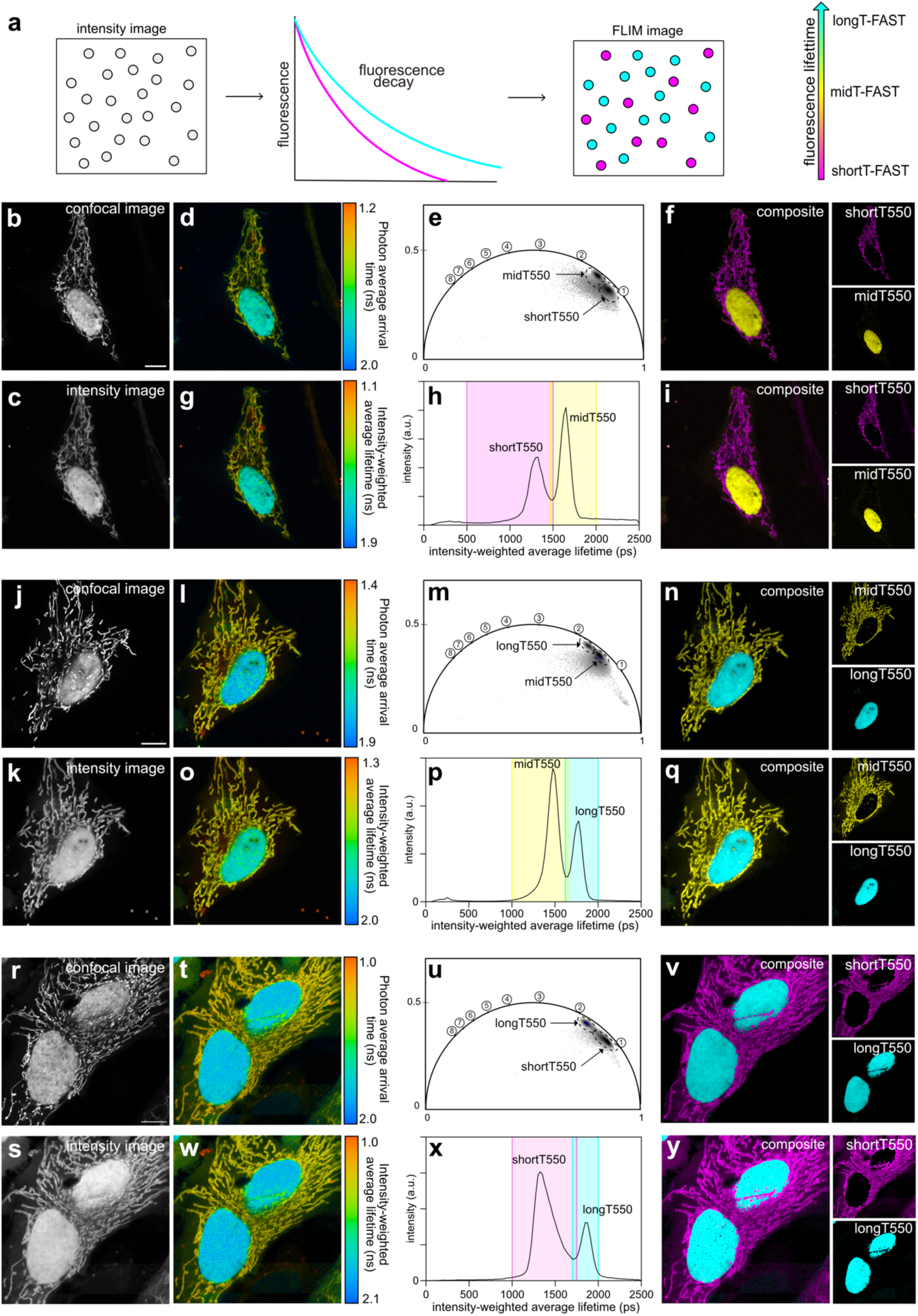
Multiplexed imaging of fluorescence lifetime modulating tags labeled with HBR-2,5DM. (a) Principle of lifetime-based separation of two FAST:fluorogen assemblies within the same sample. (**b-y**) Combinations of mitochondrial shortT550 and nuclear midT550 (**b-i**), mitochondrial midT550 and nuclear longT550 (**j-q**) and mitochondrial shortT550 and nuclear longT550 (**r-y**). For each pair, the variants were expressed in HeLa cells. For each pair, are shown (**b,j,r**) the confocal image, (**c,k,s**) the intensity image, (**d,l,t**) the photon average arrival time image, and (**g,o,w**) the intensity weighted average lifetime coded image (fit with double exponential model). (**e,m,u**) Phasor plot analysis. Each cluster corresponds to a variant (time in ns is shown on the universal circle). The clusters used for separation are circled. (**f,n,v**) Composite image resulting from the clustering shown on the phasor plot (shortT-FAST is in magenta, midT-FAST in yellow, longT-FAST in cyan). (**h,p,x**) Intensity-weighted average lifetime distribution of the sample. Each peak corresponds to a variant. In color, are shown the two windows of lifetimes used for separation. (**i,q,y**) Composite image resulting from the fitting-based separation (shortT-FAST is in magenta, midT-FAST in yellow, longT-FAST in cyan). Representative results of > 16 cells from at least three biological replicates. Scale bars, 10 µm. (**b,j,r**) Excitation 488 nm / detection window 500-600 nm.

**Fig. 3.**
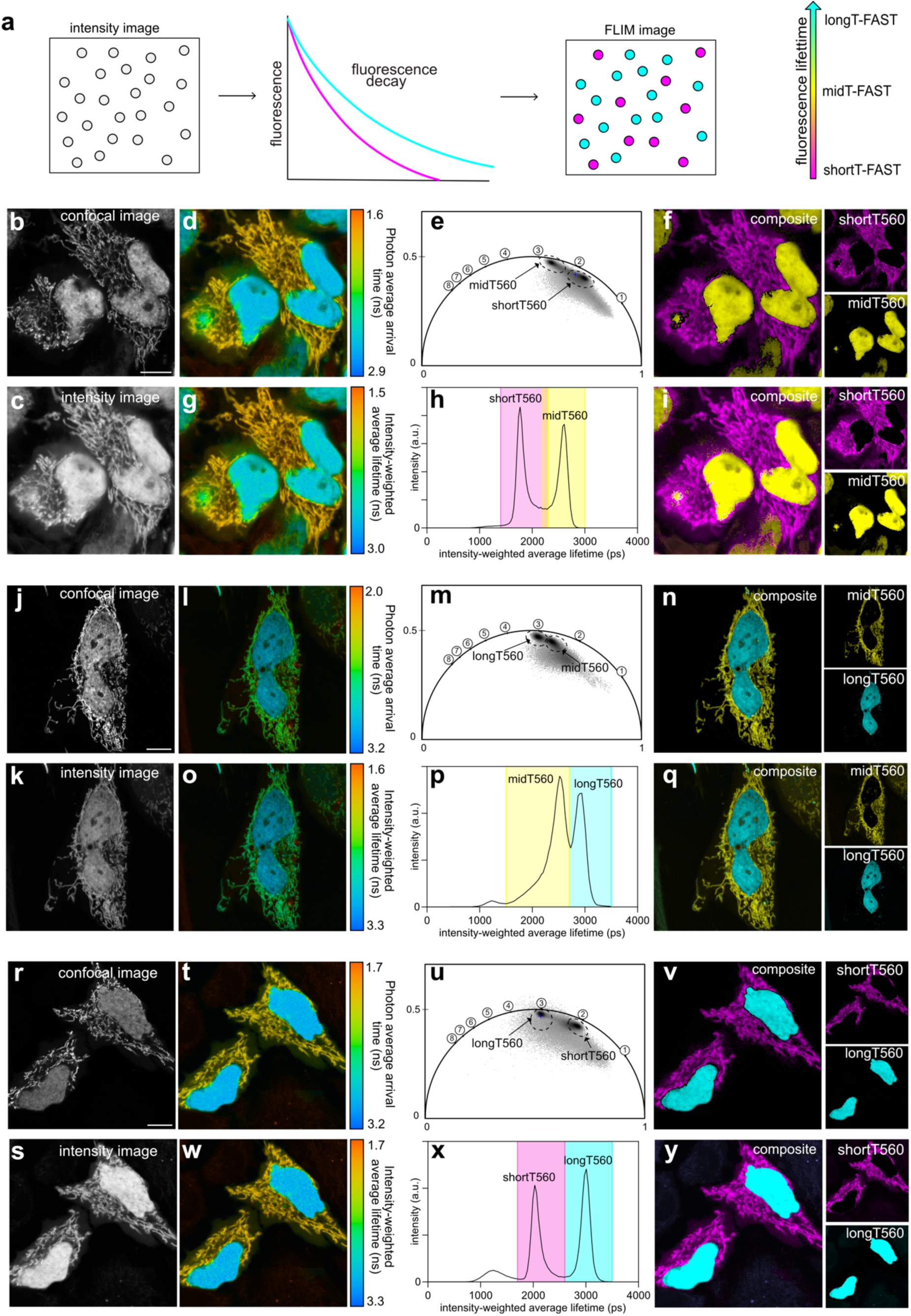
Multiplexed imaging of fluorescence lifetime modulating tags labeled with HBR-3,5DM. (a) Principle of lifetime-based separation of two FAST:fluorogen assemblies within the same sample. (**b-y**) Combinations of mitochondrial shortT560 and nuclear midT560 (**b-i**), mitochondrial midT560 and nuclear longT560 (**j-q**) and mitochondrial shortT560 and nuclear longT560 (**r-y**). For each pair, the variants were expressed in HeLa cells. For each pair, are shown (**b,j,r**) the confocal image, (**c,k,s**) the intensity image, (**d,l,t**) the photon average arrival time image, and (**g,o,w**) the intensity weighted average lifetime coded image (fit with double exponential model). (**e,m,u**) Phasor plot analysis. Each cluster corresponds to a variant (time in ns is shown on the universal circle). The clusters used for separation are circled. (**f,n,v**) Composite image resulting from the clustering shown on the phasor plot (shortT-FAST is in magenta, midT-FAST in yellow, longT-FAST in cyan). (**h,p,x**) Intensity-weighted average lifetime distribution of the sample. Each peak corresponds to a variant. In color, are shown the two windows of lifetimes used for separation. (**i,q,y**) Composite image resulting from the fitting-based separation (shortT-FAST is in magenta, midT-FAST in yellow, longT-FAST in cyan). Representative results of over >13 cells from at least three biological replicates. Scale bars, 10 µm. (**b**) Excitation 488 nm / detection window 517-600 nm. (**j,r**) Excitation 488 / detection window 517-570 nm

Based on the analysis of their fluorescence lifetimes, we reasoned that it should be possible to separate shortT550, midT550 and longT550 in the same sample. To demonstrate lifetime multiplexing of these three variants, we first co-cultured populations of HEK293T mammalian cells expressing respectively H2B–shortT-FAST, H2B–midT-FAST and H2B– longT-FAST and imaged them in presence of HBR-2,5DM by spectral confocal microscopy and FLIM (**Fig. 4a**). While spectral confocal microscopy did not allow separation of shortT550, midT550 and longT550 because of their identical spectral properties (**Fig. 4b-c**), efficient discrimination was possible through FLIM, as demonstrated by the presence of three separate lifetime populations on the photon average arrival time image (**Fig. 4d**) as well as three separate clusters on the phasor plot (**Fig. 4e**), enabling the separation of the three variants (**Fig. 4f**). When fitting the fluorescence decays with biexponential model (**Fig. 4g**), three peaks in the intensity-weighted average lifetime distribution histogram were visible at T_i_ = 1.45 ns, 1.89 ns and 2.10 ns respectively, that were attributed respectively to shortT550, midT550 and longT550 (**Fig. 4h**), allowing the separation of the three variants (**Fig. 4i**).

**Fig. 4.**
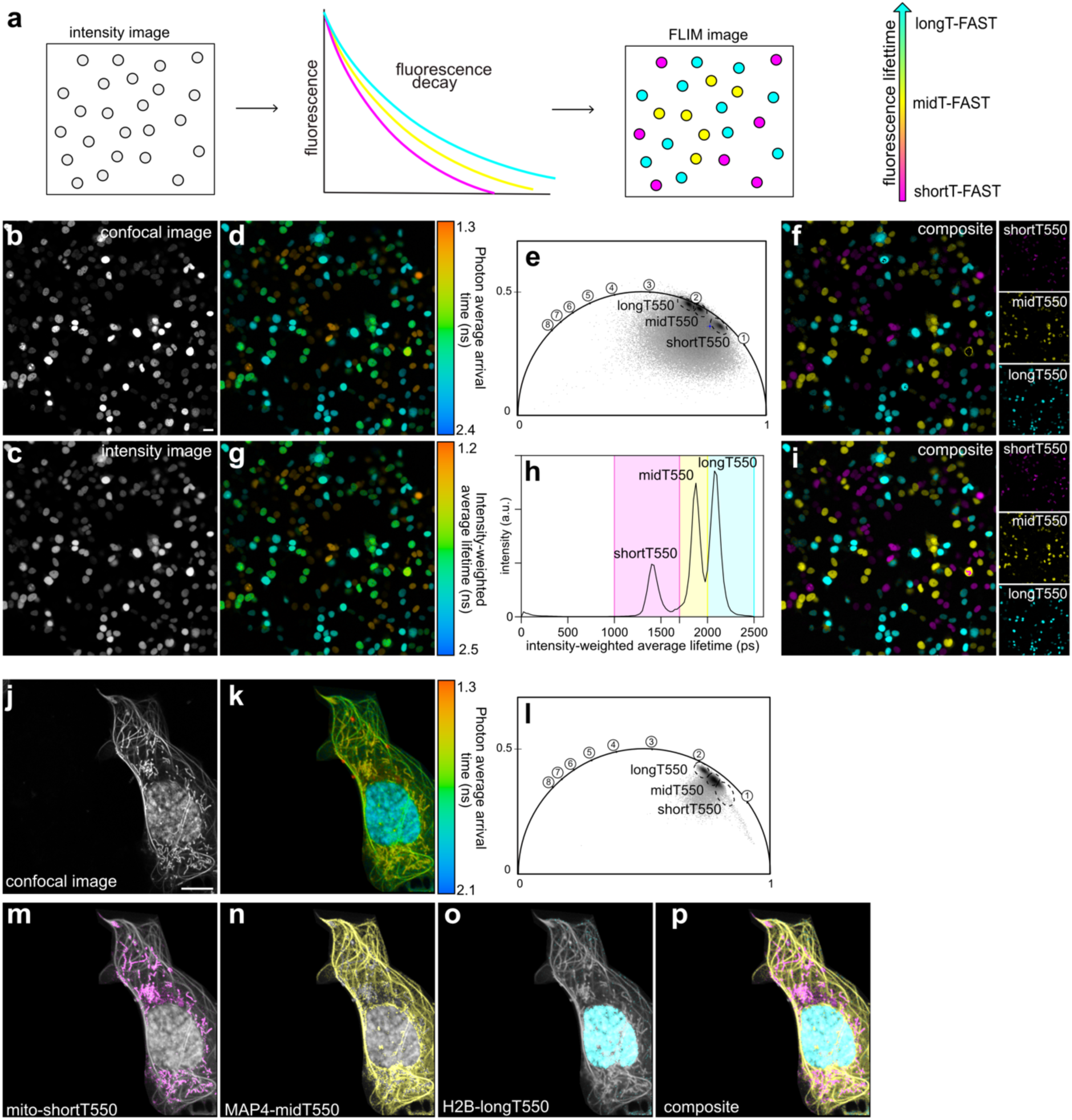
Multiplexed imaging with three fluorescence lifetime modulating tags. (a) Principle of lifetime-based separation of three FAST:fluorogen assemblies within the same sample. (**b-i**) Populations of HEK293T cells expressing either H2B−shortT-FAST or H2B−midT-FAST or H2B−longT-FAST were mixed together, labeled with 10 µM HBR-2,5DM, and imaged by confocal microscopy and FLIM. Are shown (**b**) the confocal image (Excitation 488 nm / detection window 517-600 nm), (**c**) the intensity image, (**d**) the photon average arrival time image, and (**g**) the intensity weighted average lifetime coded image (fit with double exponential model). (**e**) Phasor plot analysis. Each cluster corresponds to a variant (time in ns is shown on the universal circle). The clusters used for separation are circled. (**f**) Composite image resulting from the clustering shown on the phasor plot (shortT-FAST is in magenta, midT-FAST in yellow, longT-FAST in cyan). (**h**) Intensity-weighted average lifetime distribution of the sample. In color, are shown the three windows of lifetimes used for separation. (**i**) Composite image resulting from the fitting-based separation (shortT-FAST is in magenta, midT-FAST in yellow, longT-FAST in cyan). Scale bars, 20 µm. (**j-p**) U2OS cell expressing H2B−longT-FAST, mito−short-FAST and MAP4−midT-FAST and labeled with 10 µM HBR-2,5DM were imaged by confocal microscopy and FLIM. Are shown (**j**) the confocal image (Excitation 488 nm / detection window 517-600 nm). (**k**) the photon average arrival time image. (**l-p**) The three variants were separated using the indicated clusters on the phasor representation (**l**) and overlayed over the intensity image (**m-p**) (shortT-FAST is in magenta, midT-FAST in yellow, longT-FAST in cyan). Representative results of three independent experiments. Scale bars, 10 µm.

To further evaluate our ability to separate shortT550, midT550 and longT550, we co-expressed in U2OS shortT-FAST, midT-FAST and longT-FAST, respectively in fusion with mitochondrial targeting sequence (mito) from the subunit VIII of human cytochrome C oxidase, microtubule-associated protein (MAP)4 and H2B, and labeled them with HBR-2,5DM (**Fig. 4j,k**). Phasor plot-based identification of clusters allowed the separation of the three reporters and their visualization in the expected localization (**Fig 4l-p**). In this context, the phasor plot approach facilitated the separation of the three reporters using the positions of the individual clusters.

To further push fluorescence lifetime multiplexing in the green channel, we imaged shortT550, midT550 and longT550 together with the enhanced green fluorescent protein (EGFP) using the co-culture experiment described above. Since the reported fluorescence lifetime of EGFP is longer than that of longT550 (T_i_ = 2.6 ns, τιT_i_ = 0.5 ns)^5^, we reasoned that separation of four variants in the green channel would be possible. To unmistakably distinguish EGFP from the FAST variants, we expressed it in the cytosol (**Supplementary** Figure 9a,d). The phasor plot of the corresponding sample displayed four separated clusters (**Supplementary** Figure 9b,c), and the longest lifetime corresponded to the cells with cytosolic fluorescence, in agreement with EGFP localization. The intensity-weighted average lifetime distribution in fields of view comprising cells expressing the four different reporters displayed four separated peaks, with maxima at T_i_ = 1.26 ns, 1.64 ns, 1.82 ns and 2.08 ns, corresponding respectively to shortT550, midT550, longT550 and EGFP (**Supplementary** Figure 9e,f).

This set of experiments allowed us to demonstrate the possibility of achieving lifetime multiplexing of three different FAST variants with identical spectral properties in living cells.

### Fluorescence lifetime multiplexing in zebrafish larvae

Next, we demonstrated our ability to achieve fluorescence lifetime multiplexing of shortT-FAST, midT-FAST and longT-FAST in zebrafish larvae as model of multicellular organisms. Zebrafish is a common biological model to study a variety of biological processes such as development, inflammation, cell proliferation and metastasis^15,16^. In this context, cancer cells expressing fluorescent reporters can be injected in zebrafish embryo and larvae and their interaction with their environment as well as their proliferation can be assessed through fluorescence microscopy. To demonstrate fluorescence lifetime multiplexing of shortT550, midT550 and longT550 in a single spectral channel in zebrafish larvae, we injected a mix of three HEK293T cell populations expressing respectively H2B–shortT-FAST, H2B–midT-FAST, and H2B–longT-FAST (**Fig. 5a**) in the developing brain. Two days later, larvae were imaged after treatment with HBR-2,5DM. Confocal imaging of the larvae shows clusters of green-emitting cells near the injection point (**Fig. 5b-c**). Although some structures of the zebrafish larvae displayed some autofluorescence, their fluorescence lifetime signal did not interfere with those of the different reporters. The fluorescence lifetimes of each variant were also assessed independently through injection of the individual cell populations (**Supplementary** Figure 10) and were consistent with those measured previously in cultured mammalian cells (1.45 ns for shortT550, 1.70 ns for midT550 and 1.93 ns for longT550, **Supplementary** Figure 10d,h**,l**). Fluorescence lifetime multiplexing allowed us to discriminate the three fluorescence lifetime modulating tags in the zebrafish larvae, as demonstrated by the presence of three fluorescence lifetime signatures distinguishable in the phasor plot (**Fig. 5e-f**) as well as in the intensity-weighted average lifetime distribution (**Fig. 5g-i**). This set of experiments allowed us to demonstrate the possibility of achieving fluorescence lifetime multiplexing of three targets in a single spectral channel in a multicellular organism.

**Fig. 5.**
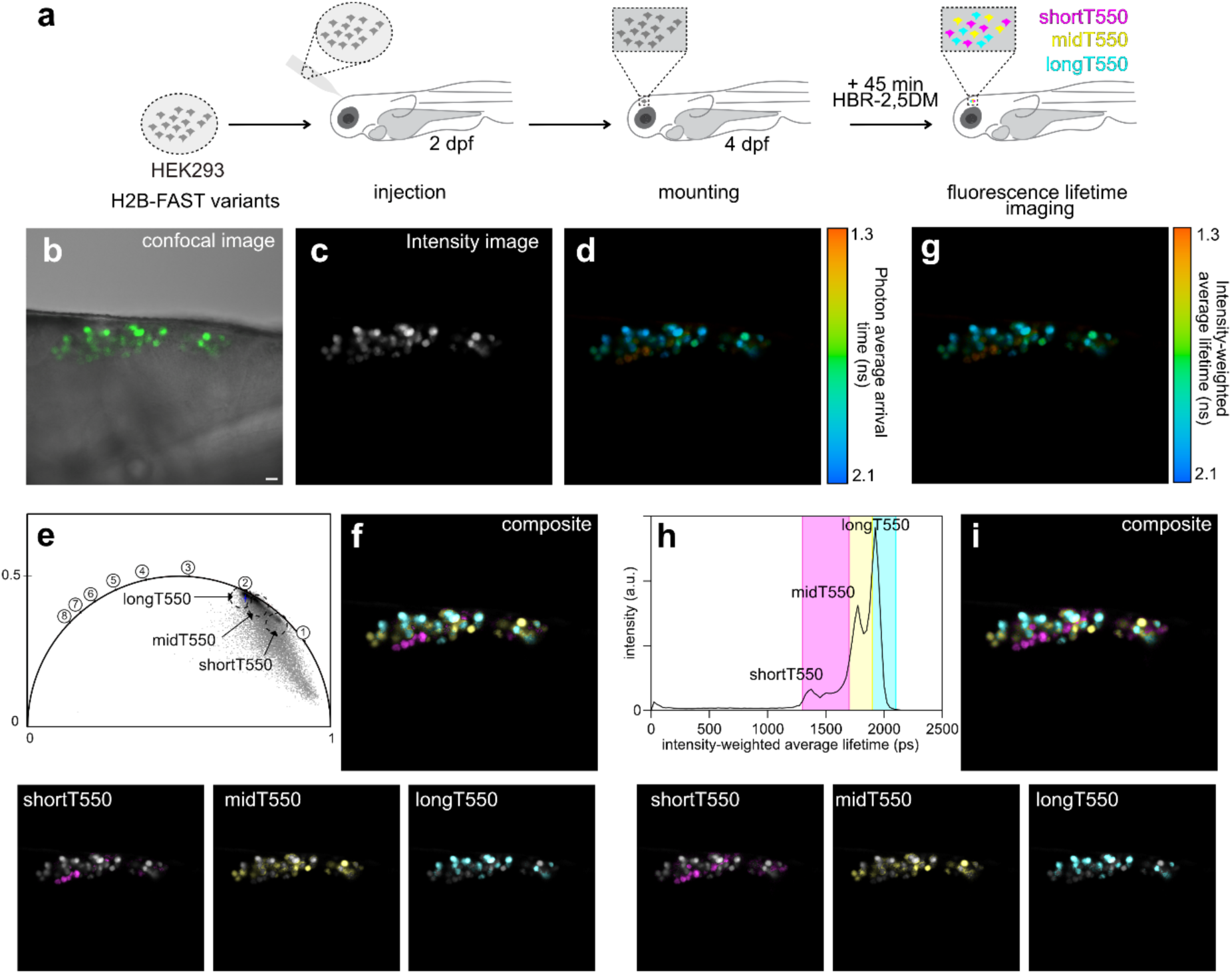
FLIM multiplexing in live zebrafish larvae. (**a**) Three populations of mammalian HEK293T were transfected with plasmid encoding either H2B-shortT-FAST, H2B-midT-FAST and H2B-longT-FAST. After 24h, they were mixed together and they were injected near the developing brain of 2 dpf zebrafish larvae. Larvae were imaged at 4 dpf after 45 min incubation with HBR-2,5DM. Are shown (**b**) the confocal image (Excitation 488 nm/ detection window 508-570 nm), (**c**) the intensity image, (**d**) the photon average arrival time image, and (**g**) the intensity weighted average lifetime coded image (fit with double exponential model). (**e**) Phasor plot analysis (time in ns is shown on the universal circle). The clusters used for separation are circled. (**f**) Composite image resulting from the clustering shown on the phasor plot. Separate channels are overlayed on the intensity channel (shortT-FAST is in magenta, midT-FAST in yellow, longT-FAST in cyan). (**h**) Intensity-weighted average lifetime distribution of the sample. In color, are shown the three windows of lifetimes used for separation. (**i**) Composite image resulting from the fitting-based separation. Separate channels are overlayed on the intensity channel (shortT-FAST is in magenta, midT-FAST in yellow, longT-FAST in cyan). Scale bars, 20 µm. Representative results of three independent experiments.

### Fluorescence lifetime multiplexing combined with spectral multiplexing

Finally, we further pushed multiplexing by combining fluorescence lifetime multiplexing with spectral multiplexing. We combined shortT550, midT550 and longT550 in the green channel, together with cyan, red and near-infrared (NIR) fluorescent reporters. For this purpose, we co-cultured mammalian cells expressing respectively ECFP (cyan channel), shortT-FAST, midT-FAST and longT-FAST (green channel), mCherry (red channel) and emIRFP670 (NIR channel) and labeled the cells with HBR-2,5DM (**Fig. 6a**). Confocal imaging allowed the spectral discrimination of ECFP, mCherry and emIRFP670 in addition to a population of green-emitting cells (**Fig. 6b-f**), and we used FLIM to separate the green fluorescence emitting reporters. Similarly to previous multiplexing experiments, fitting approach (**Fig. 6g-k**) and phasor plot (**Fig. 6l**) allowed the identification of three lifetime signatures, in agreement with the different lifetimes of shortT550, midT550 and longT550. By combining spectral and lifetime imaging, we were able to image and distinguish up to six different fluorescent reporters (**Fig. 6m**).

**Fig. 6.**
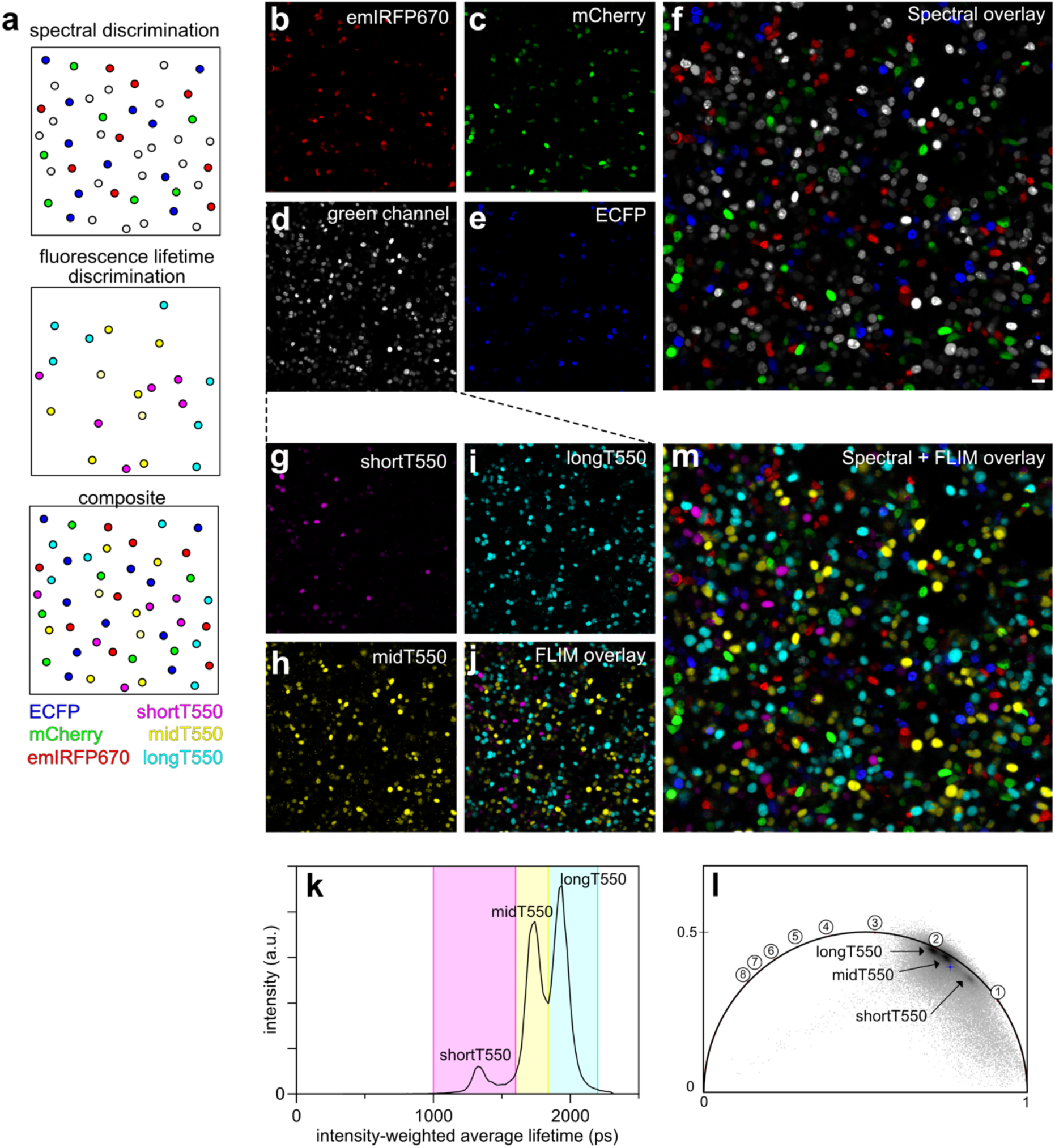
Spectral and lifetime multiplexing in live cells. (**a**) Principle of combined spectral and lifetime-based separation of six fluorescent reporters. Confocal micrographs of HEK293T cells expressing H2B-emiRFP670 (Excitation 639 nm / detection window 650-757 nm) **(b),** H2B-mCherry (Excitation 561 nm / detection window 600-632 nm) **(c),** H2B-shortT-FAST, H2B-midT-FAST, and H2B-longT-FAST (Excitation 488 nm / detection window 508-570 nm) **(d)**, and H2B-ECFP (Excitation 445 nm / detection window 455-473 nm) **(e)** in addition to spectral overlay (**f**). Scale bars, 20 µm. FLIM was then used to discriminate reporters in the green channel based on their intensity-weighted average lifetime distribution **(g-k).** The phasor plot in the green channel is also shown **(l)** (time in ns is shown on the universal circle)**. (m)** spectral and lifetime overlay. Representative results of three independent experiments. Scale bar, 20 µm.

## DISCUSSION

In this work, we leveraged a collection of FAST variants with broad sequence diversity to identify fluorescence lifetime modulating tags allowing multiplexed imaging in a single spectral channel. We screened the fluorescence lifetimes of twelve different variants with three prototypical fluorogens forming green-yellow fluorescent assemblies. Comprehensive characterization of greenFAST, oFAST and TsiA-FAST, renamed shortT-FAST, midT-FAST and longT-FAST, respectively, showed that these three variants formed fluorescent assemblies with HBR-2,5DM and HBR-3,5DM that display distinct fluorescence lifetime signatures, allowing their separation by FLIM although their spectral properties are identical. These differences in fluorescence lifetime are not due to differences in their fluorescence quantum yields, but rather originate from different non-radiative deexcitation behaviors. Our results suggest that the fluorescence lifetimes of such fluorogenic reporters can be tuned through engineering of the binding pocket of the fluorophore, opening interesting prospects for engineering variants with more substantial differences in lifetime: while the screening strategy previously used during the directed evolution process of FAST proteins focused more on selecting the tightest and brightest assemblies, an additional screening based on fluorescence lifetime could enable in the future the identification of variants displaying more significant differences in lifetime, and allow multiplexing of more targets. Among the three variants highlighted in this study, longT-FAST (a.k.a TsiA-FAST) was not obtained through directed evolution like the two others, but by exploring proteins with homology relationship. While this homology-based protein engineering strategy allowed the generation of variants with sequence similarity of 70-78 %, exploring the use of homologs with lower similarity may allow the generation of novel FAST variants with different photophysical behaviors and thus different fluorescence lifetime signatures.

To demonstrate the possibility of achieving fluorescence lifetime multiplexing, we performed pairwise combinations of shortT-FAST, midT-FAST and longT-FAST with either HBR-2,5DM or HBR-3,5DM, and showed that, for a given fluorogen, proper discrimination of each possible pair could be achieved. With HBR-2,5DM, shortT550, midT550 and longT550 could be combined in the same experiment for fluorescence lifetime multiplexing of three targets in a single spectral channel as demonstrated in cultured mammalian cells. We leveraged this feature in zebrafish larvae through injection of a mixture of three mammalian cell populations, each expressing a different fluorescence lifetime modulating tag, and successfully distinguished them via their different lifetime signatures.

One interesting feature of fluorescence lifetime multiplexing is the possibility to distinguish variants with different lifetime signatures in one spectral channel, freeing up the other spectral channels for observing additional biological targets^20,21^. The fluorescence lifetime modulating tags identified in this study are green fluorescent. We demonstrated that spectral multiplexing and lifetime multiplexing could be combined to discriminate up to six different fluorescent reporters, opening prospects for monitoring as many different processes simultaneously.

The identification of fluorescence lifetime modulating tags for multiplexing enables us to envision the use of these systems as reporters for the design of lifetime responsive biosensors through coupling with recognition domains of specific analytes and biomolecules of interest. We believe fluorescence lifetime multiplexing using chemogenetic fluorescence lifetime modulating tags open exciting prospects for biological imaging: combined with spectral multiplexing, it opens countless opportunities for observing several biological targets simultaneously.

## METHODS

### General

Commercially available reagents were used as obtained. The synthesis of HBR-2,5DM^22^, HBR-3,5DM^22^ and HMBR^8^ was previously reported. These fluorogens are commercially available from the Twinkle Factory under the name match_550_, ^TF^Amber and ^TF^Lime.

### Biology

The presented research complies with all relevant ethical regulations.

### General

Synthetic oligonucleotides used for cloning were purchased from Integrated DNA Technology. PCR reactions were performed with Q5 polymerase (New England Biolabs) in the buffer provided. PCR products were purified using QIAquick PCR purification kit (QIAGEN). DNase I, T4 ligase, fusion polymerase, Taq ligase and Taq exonuclease were purchased from New England Biolabs and used with accompanying buffers and according to the manufacturer’s protocols. Isothermal assemblies (Gibson Assembly) were performed using a homemade mix prepared according to previously described protocols^17^. Small-scale isolation of plasmid DNA was conducted using a QIAprep miniprep kit (QIAGEN) from 2 mL overnight bacterial culture supplemented with appropriate antibiotics. Large-scale isolation of plasmid DNA was conducted using the QIAprep maxiprep kit (QIAGEN) from 150 mL overnight bacterial culture supplemented with appropriate antibiotics. All plasmid sequences were confirmed by Sanger sequencing with appropriate sequencing primers (GATC Biotech).

### Cloning

The plasmids used in this study have been generated using isothermal Gibson Assembly or restriction enzymes cloning.

The plasmids pAG641, pAG261, pAG382 and pAG645 enabling the bacterial expression of 6 ×His–TEVcs–pFAST^9^, 6 ×His–TEVcs–greenFAST^13^, 6 ×His–TEVcs–TsiA-FAST^14^ and 6 ×His–TEVcs–oFAST^9^ were previously described.

The construction of plasmids pAG109 pAG374, pAG472, pAG657, pAG658 and pAG659 allowing the mammalian expression of H2B-FAST^8^, H2B-greenFAST^13^, H2B-iFAST^12^, H2B-pFAST^9^, H2B-tFAST^9^ and H2B-oFAST^9^ was previously described. The construction of plasmids pAG372 and pAG673 allowing the expression of mito-greenFAST^13^ and mito-oFAST^9^ was previously described. The plasmids pAG765, pAG766, pAG767, pAG768, pAG769 and pAG770 allowing the expression of cMyc-HboL-FAST-IRES-mTurquoise2, cMyc-HspG-FAST-IRES-mTurquoise2, cMyc-RspA-FAST-IRES-mTurquoise2, cMyc-Ilo-FAST-IRES-mTurquoise2, cMyc-TsiA-FAST-IRES-mTurquoise2, cMyc-Rsa-FAST-IRES-mTurquoise2 were previously described^14^. The plasmids pAG1367 and pAG1448 allowing the mammalian expression of H2B-emIRFP670-cMyc and H2B-TsiA-FAST-cMyc respectively were constructed by replacing the sequence of pFAST by the sequence of TsiA-FAST in the plasmid pAG657 allowing the expression of H2B-pFAST-cMyc. The plasmids pAG324 and pAG1247 allowing the expression of H2B-mCherry^18^ and H2B-ECFP-pFAST_1-114_^19^ were previously described. The plasmid pAG1052 allowing the expression of FRB-EGFP was obtained by replacing the sequence of frFAST_1-114_ by the sequence of EGFP in the plasmid pAG499 allowing the expression of CMV-cMyc-FRB-N-frFAST^18^

### Cell culture

HeLa cells (ATCC CRM-CCL2) were cultured in minimal essential medium supplemented with phenol red, Glutamax I, 1 mM of sodium pyruvate, 1% (vol/vol) of non-essential amino acids, 10% (vol/vol) fetal calf serum (FCS) and 1% (vol/vol) penicillin– streptomycin at 37 °C in a 5% CO_2_ atmosphere. HEK293T (ATCC CRL-3216) cells were cultured in Dulbecco’s modified Eagle medium (DMEM) supplemented with phenol red, 10% (vol/vol) FCS and 1% (vol/vol) penicillin–streptomycin at 37 °C in a 5% CO_2_ atmosphere. U2OS cells (ATCC HTB-96) were cultured in McCoy’s medium supplemented with phenol red and 10% (vol/vol) FCS and 1% (vol/vol) penicillin–streptomycin at 37 °C in a 5% CO_2_ atmosphere. For imaging, cells were seeded in μDish IBIDI (Biovalley) coated with poly-L-lysine. Cells were transiently transfected using Genejuice (Merck) according to the manufacturer’s protocols for 24 h before imaging. Cells were washed with Dulbecco’s PBS (DPBS) and treated with DMEM (without serum and phenol red) supplemented with the compounds at the indicated concentration.

### Protein expression in bacteria and purification

Plasmids were transformed in BL21(DE3) competent *Escherichia coli* (New England Biolabs) or Rosetta(DE3)pLysS *E. coli* (Merck). Cells were grown at 37 °C in lysogeny broth medium supplemented with 50 μg .mL^−1^ kanamycin (and 34 μg.mL^−1^ of chloramphenicol for Rosetta) to OD_600nm_ 0.6. Expression was induced overnight at 16 °C by adding isopropyl β-D-1-thiogalactopyranoside (IPTG) to a final concentration of 1 mM. Cells were collected by centrifugation (4,300 × g for 20 min at 4 °C) and frozen. For purification, the cell pellet was resuspended in lysis buffer (PBS supplemented with 2.5 mM MgCl_2_, 1 mM of protease inhibitor phenylmethanesulfonyl fluoride and 0.025 mg.mL^−1^ DNase, pH 7.4) and sonicated (5 min, 20% of amplitude) on ice. The lysate was incubated for 2 h on ice to allow DNA digestion by DNase. Cellular fragments were removed by centrifugation (9,000 × g for 1 h at 4 °C). The supernatant was incubated overnight at 4 °C by gentle agitation with pre-washed Ni-NTA agarose beads in PBS buffer complemented with 20 mM of imidazole. Beads were washed with ten volumes of PBS complemented with 20 mM of imidazole and with five volumes of PBS complemented with 40 mM of imidazole. His-tagged proteins were eluted with five volumes of PBS complemented with 0.5 M of imidazole. The buffer was exchanged to PBS (0.05 M phosphate buffer and 0.150 M NaCl) using PD-10 desalting columns or Midi-Trap G-25 (GE Healthcare). The purity of the proteins was evaluated using SDS–PAGE electrophoresis stained with Coomassie blue.

### Physicochemical measurements

Steady-state UV-Vis and fluorescence spectra were recorded at 25 °C on a Spark 10 M (Tecan). Data were processed using GraphPad Prism v.10.0.3. Fluorescence quantum yield measurements were determined in 96-well plates using either FAST:HMBR or pFAST:HBR-3,5DM as a reference. Solutions of 40 µM of proteins were used, in which the fluorogen was diluted to the right concentration, usually from 6 µM to 0.375 µM, allowing > 99% of complex formation. Absorption coefficients were determined directly by the previous experiments after determination of the optical path length. Thermodynamic dissociation constants were determined by titration experiments in which we measured the fluorescence of the fluorescent assembly at various fluorogen concentrations using a Spark 10M plate reader (Tecan) and fitting data in Prism 9 to a one-site specific binding model.

### Fluorescence microscopy

The confocal micrographs were acquired on a Zeiss LSM 980 Laser Scanning Microscope equipped with a plan apochromat 20 × dry (NA 0.8) objective and with a plan apochromat 63× /1.4 NA oil immersion objective ZEN software were used to collect the confocal data. Fiji was used to analyze the confocal data.

## Fluorescence lifetime imaging

### General

FLIM images were acquired on a Zeiss LSM 980 confocal system coupled with Becker and Hickl FLIM module and equipped with a 20 × dry (NA 0.8) objective or a 63 × oil immersion (NA 1.4) objective. A 488 nm pulsed diode laser (50 MHz) for the excitation, a green large band bloc filter (DBS 450 + 538; DBP 467/24 + 598/110) and a HPM-100-40-ZEISS Hybrid GaAsP Photodetector were used. Acquisitions were performed at 37°C for mammalian cells and 30°C for zebrafish. All FLIM analysis was performed on the software SPCImage of Becker and Hickl^23^.

### Single variants characterization

For each variant, at least six fields of view using a 20 × objective from two biological replicates were acquired, first through confocal imaging then through lifetime imaging. For lifetime determination, regions of interest were drawn around the nuclei, and the corresponding fluorescence decays were fitted with mono– or bi-exponential models (using maximum likelihood estimation (MLE) for better accuracy). For each fitting model, the fluorescence lifetime components were reported, and the intensity-weighted average lifetime was computed. For each FAST:fluorogen assembly, > 70 cells from two biological replicates were analyzed this way. The mean of intensity-weighted average fluorescence lifetime was used as a parameter to characterize the fluorescent assemblies.

### Fluorescence lifetime multiplexing

For each pair, confocal micrographs using 63 × objective were acquired prior to lifetime imaging to ensure optimal focus and thus better data quality. After lifetime acquisition, separation of the two variants was performed systematically through:

(i) Calculation of the first moment: this gives a distribution of average photon arrival time. Variants that can be separated based on their lifetimes appear in different colors.
(ii) Fitting of the fluorescence decays with bi-exponential model and plotting of the corresponding intensity-weighted lifetime distribution through the whole image. Variants that can be separated will result in two distinct peaks in the resulting histogram and selection of lifetime windows around the peaks enables separation of the two variants.
(iii) After drawing a region of interest around the cell to analyze, the corresponding phasor plot can be displayed, and pixels with similar lifetimes form a cluster. Two distinct clusters are visible when the variants can be separated. Pixels forming each cluster can be identified in the lifetime image and thus separated.

### Zebrafish experiments

Adult zebrafish (*Danio rerio*) were kept at around 27-29°C on a 14 hr-light: 10 hr-dark cycle and fed twice daily. Natural crosses obtained fertilized eggs which were raised at 28°C in Volvic water. Experiments were performed using the standard nacre strain. Developmental stages were determined and indicated as days post fertilization (dpf). The animal facility obtained permission from the French Ministry of Agriculture (agreement No. D-75-05-32.) and all animal procedures were performed in accordance with French animal welfare guidelines. HEK293T were seeded at 500,000 cells/mL concentration in 25 cm^2^ flasks and transfected after 24h using GeneJuice (Merck) according to manufacturer’s guidelines. After 24h, cells were harvested at 10,000,000 cells/mL concentration in serum free DMEM. Cell suspension was loaded into a borosilicate glass needle pulled by a Flaming/Brown micropipette puller (Narishige, Japan, PN-30). 5∼10 nanoliters suspension were implanted into anesthetized (0.02% MS-222 tricaine (Sigma)) 2 dpf zebrafish larvae within the developing brain by using an electronically regulated air-pressure microinjector (FemtoJet, Eppendorf). After injection, zebrafish larvae were placed in Volvic water and examined under a stereoscopic microscope for the presence of fluorescent cells and then raised for two more days at 28°C before imaging. For imaging, living zebrafish larvae were anesthetized in MS-222 tricaine solution and embedded in a lateral orientation in low-melting agarose (0.8%). Larval zebrafish were studied before the onset of sexual differentiation and their sex can therefore not be determined.

## ACKNOWLEDGMENTS

This work has been supported by the European Research Council (ERC-2016-CoG-724705 FLUOSWITCH), the Institut Universitaire de France, and the Fondation pour la Recherche Médicale (grant FDT202304016615 to LEH). LEH thanks the École Normale Supérieure for PhD funding. The authors acknowledge the IBPS Imaging facility. Image acquisition and/or image analysis were performed at the IBPS Imaging Facility. The IBPS Imaging facility is supported by Region-Île-de-France, Sorbonne Université, CNRS and GIS-IBISA.

## AUTHOR CONTRIBUTIONS

L.E.H and A.G. designed the overall project and wrote the paper with the help of the other authors. L.E.H, F. L., C.R., M.V., S.V. and A.G. designed the experiments. L.E.H, F.L., M.A., H.B., C.R., M.V. and S.V. performed the experiments. L.E.H, F.L., C.R., M.V., S.V. and A.G. analyzed the experiments.

## DATA AVAILABILITY

The data supporting the findings of this study are available within the article and supplementary information, and are available from the corresponding authors upon reasonable request. The plasmids developed in this study will be available from Addgene.

