## Supporting information for "Multiplexed in vivo imaging with fluorescence lifetime modulating tags"

**This PDF file includes:**

Supplementary Text 1

Supplementary Figures 1-10

Supplementary Tables 1-3

**Supplementary Text 1. The FAST family**

The Fluorescence-activating and absorption-shifting tag (FAST) toolbox is a family of small chemogenetic reporters evolved from the 14-kDa photoactive yellow protein (PYP) found in the phototactic bacterium *Halorhodospira halophila*. Prototypical FAST binds hydroxybenzylidene rhodanine (HBR) derivatives[1,2] and stabilizes their fluorescent state. When they are free in solution, these so-called fluorogens dissipate energy through non-radiative de-excitation pathways. However, within the cavity of FAST, they adopt a quasi-planar conformation, and the phenolate form of the fluorogens is stabilized through hydrogen bonding with residue E46, a key residue for the modulation of spectral properties of PYP[3,4].

Since the development of prototypical FAST, the FAST family of reporters has been fairly extended through alteration of the structure of the fluorogens, as well as engineering of adequate cognate protein tags. Molecular engineering of the phenol ring of HBR scaffold allowed the design of new fluorogens displaying green to red emission in assembly with FAST, allowing to have different fluorescence emission properties with the same protein tag using either HBR-2,5DM, HBR-3,5DM and HBR-3,5DOM[2]. Rational design allowed the development of an enhanced version of FAST named iFAST, characterized by superior properties with HMBR[5]. Directed evolution later allowed the design of two orthogonal tags, greenFAST and redFAST, binding preferably HMBR and HBR-3,5DOM respectively. The characterization of the fluorescence lifetime of greenFAST:HMBR assembly highlighted an interesting behaviour, as its fluorescence lifetime was shown to be shorter than that of iFAST:HMBR, although the two assemblies have similar fluorescence quantum yields (FQY). More recently, the FAST family was extended with three new reporters: pFAST, oFAST and tFAST, obtained through a concerted strategy of directed evolution and molecular engineering of the chromophore[6]. pFAST has the advantageous feature of being a promiscuous tag, able to bind and activate the fluorescence of different fluorogens families, enabling fluorescent assemblies spanning the visible spectrum.

While the previously cited FAST variants were engineered through directed evolution, a novel strategy based on protein homology allowed recently the development of six new FAST reporters, HboL-FAST, HspG-FAST, RspA-FAST, Ilo-FAST, TsiA-FAST and Rsa-FAST[7], engineered from six homologs of PYP from *Halomonas boliviensis* LC1 (HboL), *Halomonas* sp. GFAJ-1 (HspG), *Rheinheimera* sp. A13L (RspA), *Idiomarina loihiensis* (Ilo), *Thiorhodospira sibirica* ATCC 700588 (TsiA), and *Rhodothalassium salexigens* (Rsa), displaying 70-78% sequence similarity with the original FAST sequence.

These FAST mutants constitute an interesting platform for the identification of variants which can be multiplexed based on their lifetime properties.

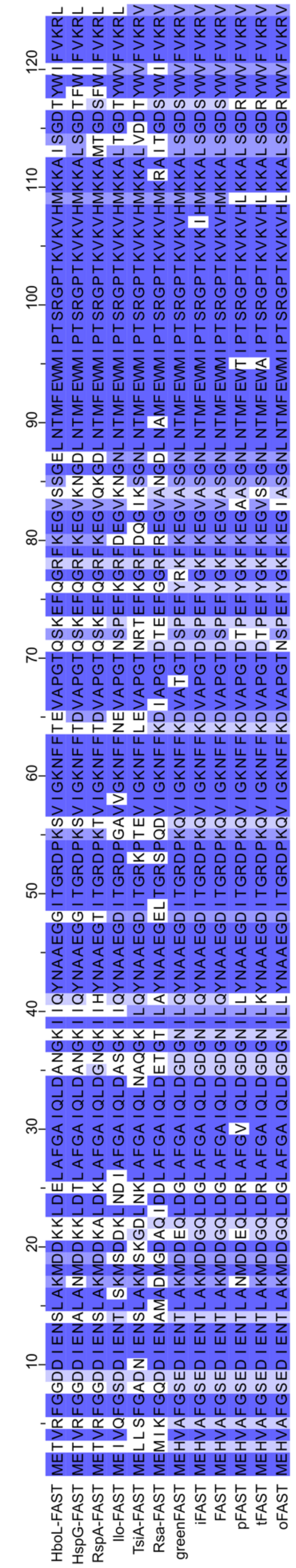


**Supplementary Figure 1. Alignment of the FAST variants used in this study.**

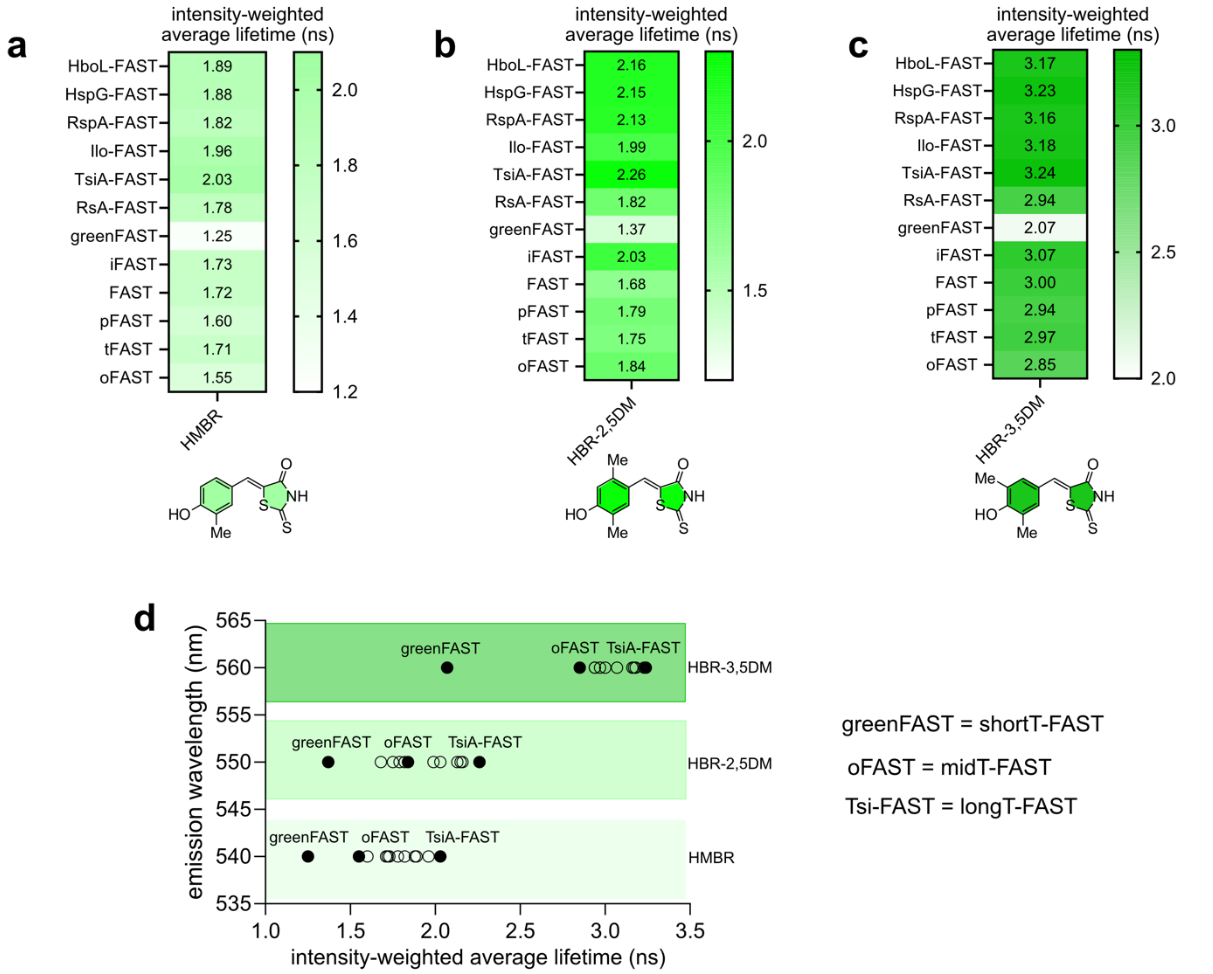


**Supplementary Figure 2. Fluorescence lifetime screening of FAST:fluorogen pairs.** Mean (n = 8-16 cells) of intensity-weighted average lifetimes of FAST variants with (**a**) HMBR, (**b**) HBR-2,5DM and (**c**) HBR-3,5DM as measured in HEK293T cells. (**d**) Emission wavelengths of the FAST:fluorogen assemblies against their intensity-weighted average lifetimes.

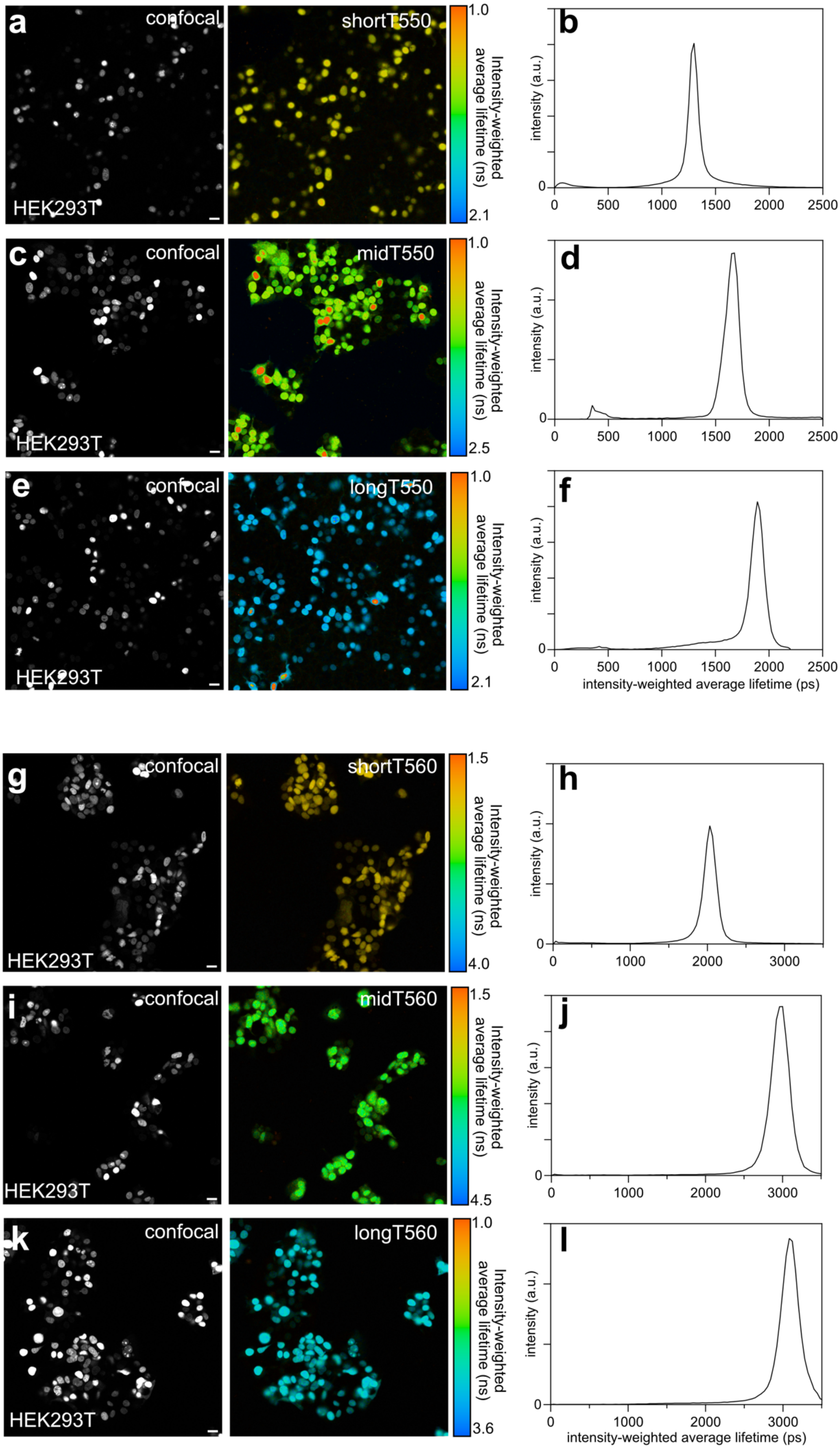

a
confocal
HEK293T
shortT550
1.0
2.1
Intensity-weighted average lifetime (ns)
b
intensity (a.u.)
c
midT550
2.5
d
e
longT550
f
intensity-weighted average lifetime (ps)
g
shortT560
1.5
4.0
h
i
midT560
4.5
j
k
longT560
3.6
l

**Supplementary Figure 3. Fluorescence lifetime characterization of shortT-FAST, midT-FAST and longT-FAST with HBR-2,5DM and HBR-3,5DM in HEK293T cells.** Confocal micrographs, equivalent intensity-weighted average lifetime images and intensity-weighted average lifetime histogram (fit with double-exponential model) of HEK293T cells expressing respectively H2B-shortT-FAST (**a,b, g,h**), H2B-midT-FAST (**c,d, i,j**) and H2B-longT-FAST (**e,f, k,l**) labeled with 10 µM HBR-2,5DM (**a-f**) or 10 µM HBR-3,5DM (**g-l**). Representative micrographs and distribution of at least six fields of view from two independent experiments. Scale bars, 20 µm. (**a,c,e,g,i,k**) Excitation wavelength 488 nm / Detection window 517-600 nm.

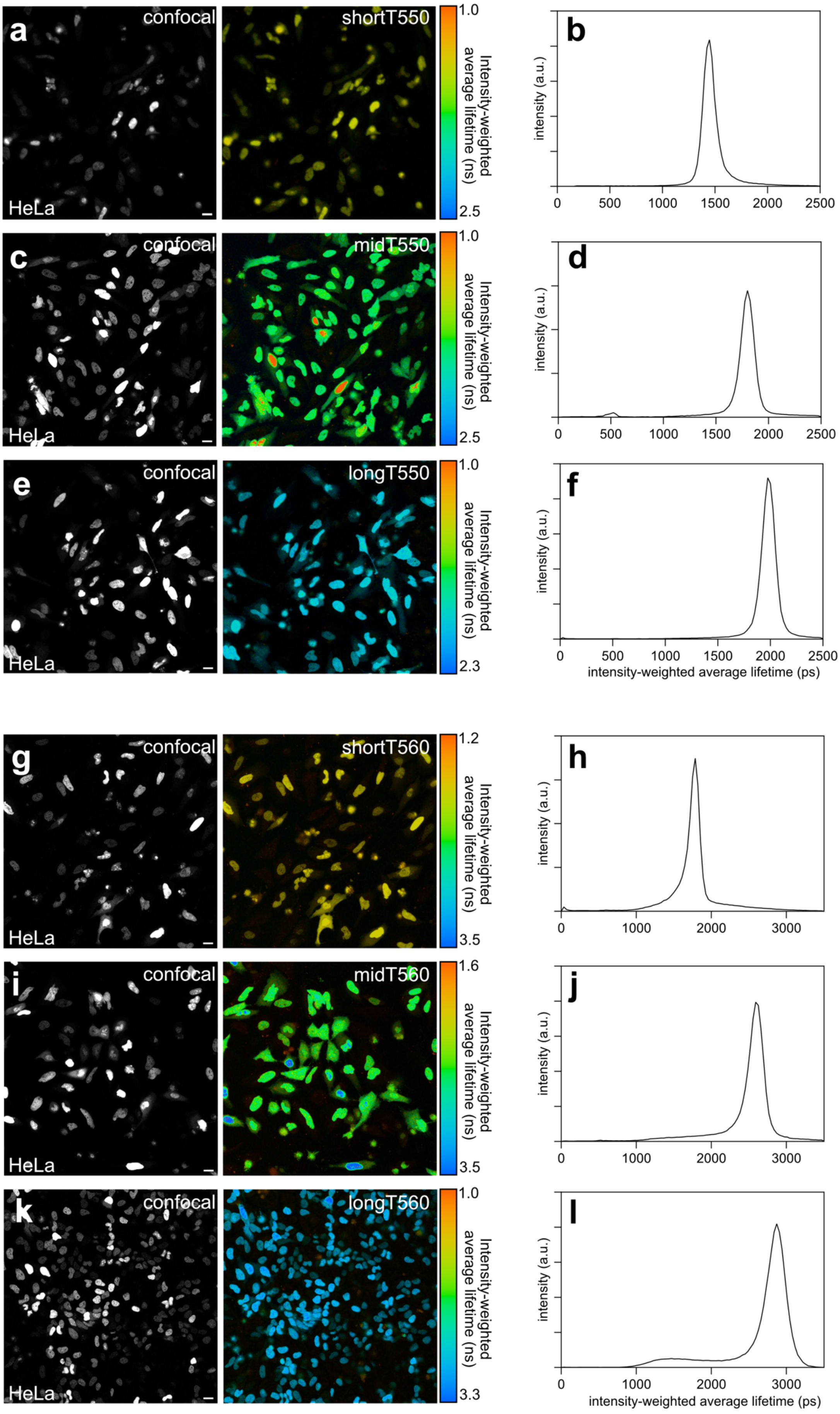

a
confocal
HeLa
shortT550
1.0
2.5
Intensity-weighted average lifetime (ns)
b
intensity (a.u.)
0
500
1000
1500
2000
2500
c
confocal
HeLa
midT550
1.0
2.5
Intensity-weighted average lifetime (ns)
d
intensity (a.u.)
e
confocal
HeLa
longT550
1.0
2.3
Intensity-weighted average lifetime (ns)
f
intensity (a.u.)
intensity-weighted average lifetime (ps)
g
confocal
HeLa
shortT560
1.2
3.5
Intensity-weighted average lifetime (ns)
h
intensity (a.u.)
0
1000
2000
3000
i
confocal
HeLa
midT560
1.6
3.5
Intensity-weighted average lifetime (ns)
j
intensity (a.u.)
k
confocal
HeLa
longT560
1.0
3.3
Intensity-weighted average lifetime (ns)
l
intensity (a.u.)
intensity-weighted average lifetime (ps)

**Supplementary Figure 4. Fluorescence lifetime characterization of shortT-FAST, midT-FAST and longT-FAST with HBR-2,5DM and HBR-3,5DM in HeLa cells.** Confocal micrographs, equivalent intensity-weighted average lifetime images and intensity-weighted average lifetime histogram (fit with double-exponential model) of HeLa cells expressing respectively H2B-shortT-FAST (**a,b, g,h**), H2B-midT-FAST (**c,d, i,j**) and H2B-longT-FAST (**e,f, k,l**) labeled with 10 µM HBR-2,5DM (**a-f**) or 10 µM HBR-3,5DM (**g-l**)**.** Representative micrographs and distribution of at least six fields of view from two independent experiments. Scale bars, 20 µm. (**a,c,e,g,i,k**) Excitation 488 nm / detection window 517-600 nm.

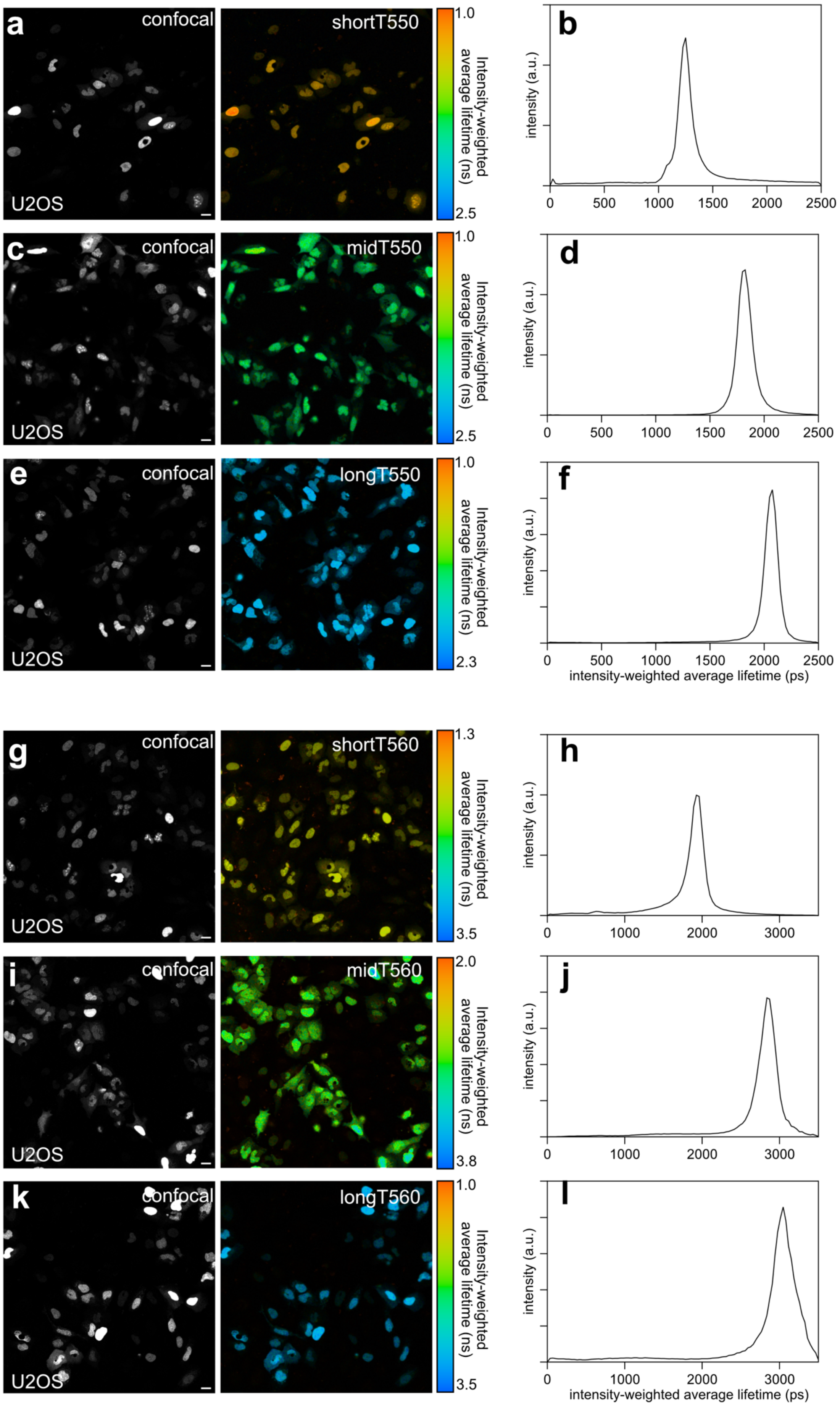
a
confocal
U2OS
shortT550
1.0
2.5
Intensity-weighted average lifetime (ns)
b
intensity (a.u.)
0
500
1000
1500
2000
2500
c
confocal
U2OS
midT550
1.0
2.5
Intensity-weighted average lifetime (ns)
d
intensity (a.u.)
0
500
1000
1500
2000
2500
e
confocal
U2OS
longT550
1.0
2.3
Intensity-weighted average lifetime (ns)
f
intensity (a.u.)
0
500
1000
1500
2000
2500
intensity-weighted average lifetime (ps)
g
confocal
U2OS
shortT560
1.3
3.5
Intensity-weighted average lifetime (ns)
h
intensity (a.u.)
0
1000
2000
3000
i
confocal
U2OS
midT560
2.0
3.8
Intensity-weighted average lifetime (ns)
j
intensity (a.u.)
0
1000
2000
3000
k
confocal
U2OS
longT560
1.0
3.5
Intensity-weighted average lifetime (ns)
l
intensity (a.u.)
0
1000
2000
3000
intensity-weighted average lifetime (ps)

**Supplementary Figure 5. Fluorescence lifetime characterization of shortT-FAST, midT-FAST and longT-FAST with HBR-2,5DM and HBR-3,5DM in U2OS cells.** Confocal micrographs, equivalent intensity-weighted average lifetime images and intensity-weighted average lifetime histogram (fit with double-exponential model) of U2OS cells expressing respectively H2B-shortT-FAST (**a-b, g-h**), H2B-midT-FAST (**c-d, i-j**) and H2B-longT-FAST (**e-f, k-l**) labeled with 10 μM HBR-2,5DM (**a-f**) or 10 μM HBR-3,5DM (**g-l**)**.** Representative micrographs and distribution of at least six fields of view from two independent experiments. Scale bars, 20 μm. (**a,c,e,g,i,k**) Excitation 488 nm / detection window 517-600 nm.

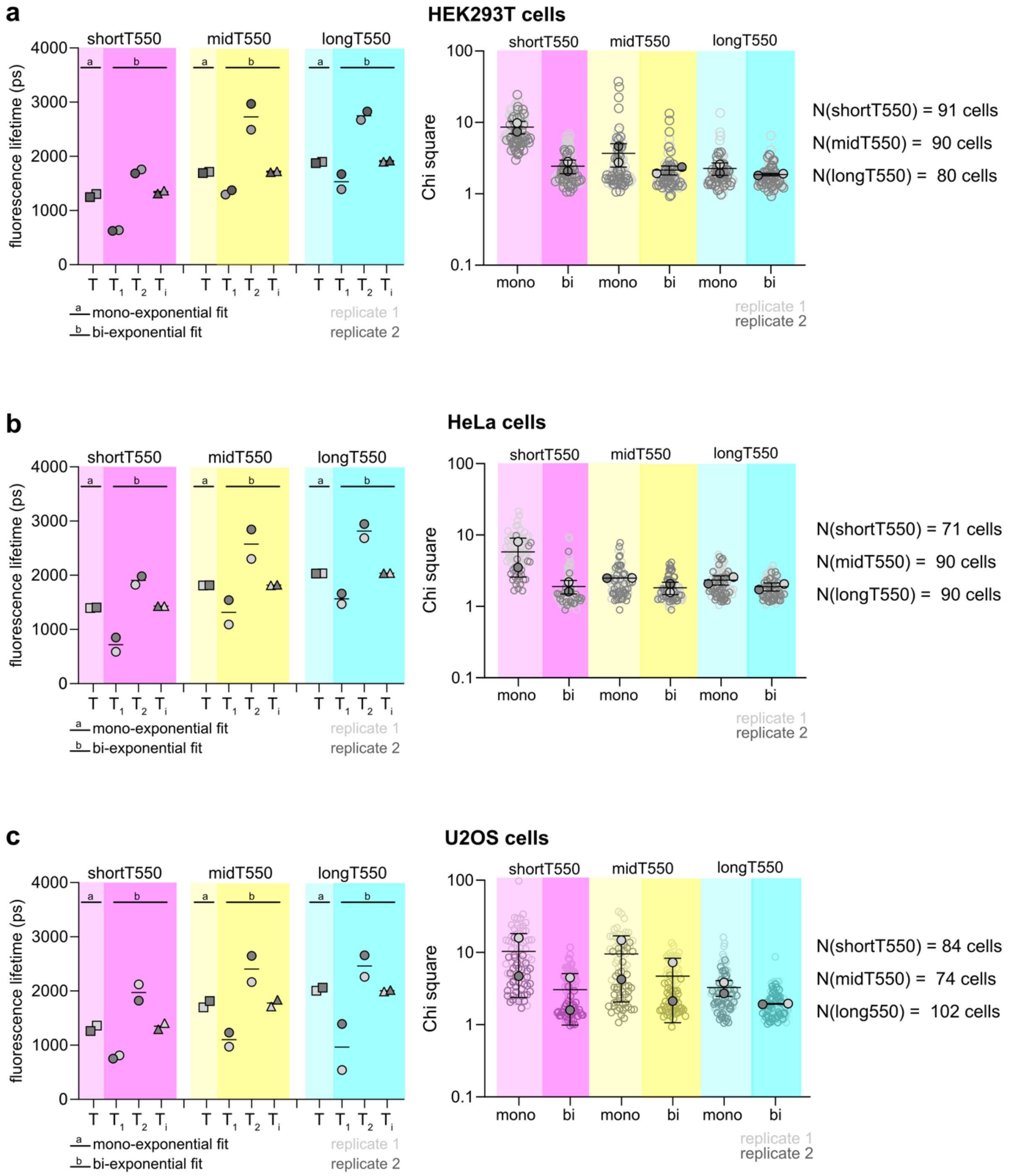


**Supplementary Figure 6. Comparison of the fitting models for the fluorescence lifetime analysis of shortT550, midT550 and longT550 in different cell lines.** Means of fluorescence lifetime T of shortT550, midT550 and longT550 extracted from fitting with single-exponential model, as well as of the two fluorescence lifetime components $T_1$ and $T_2$ and intensity-weighted average fluorescence lifetime $T_i$ from fitting with bi-exponential model in (**a**) HEK293T cells, (**b**) HeLa cells and (**c**) U2OS cells. N cells from two biological replicates were analyzed (the number N of analyzed cells is indicated on the right). The black lines represent the means of the two biological replicates. Each dot corresponds to the mean of each biological replicate (the lifetimes of individual cells are not shown). On the right is shown the distribution of goodness of fit ( reduced $\chi^2$ or chi square) of mono- and bi-exponential models. Each cell is color-coded according to the biological replicate it came from. The solid circles correspond to the mean of each biological replicate. The black line represents the mean ± SD of the two biological replicates.

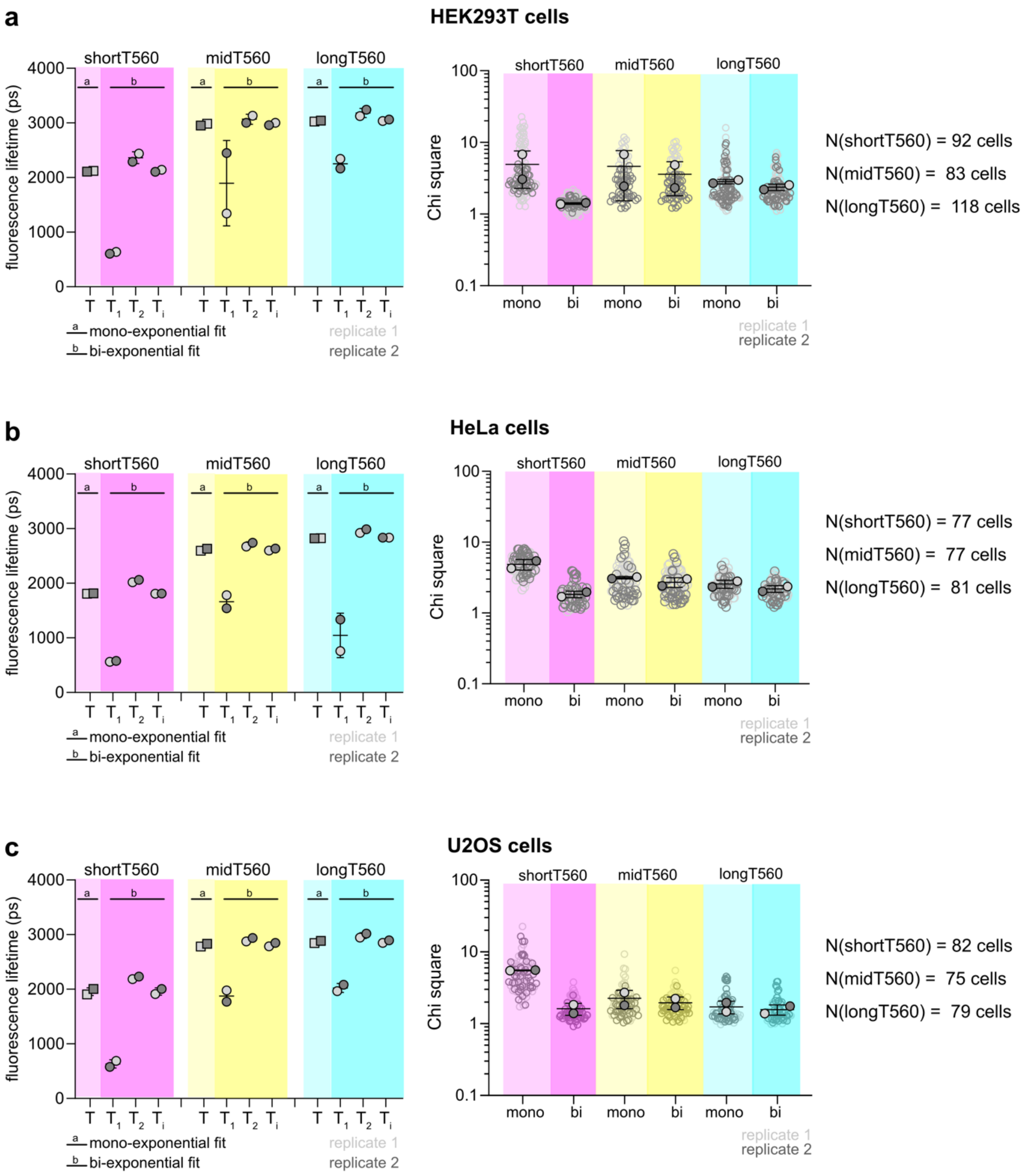


**Supplementary Figure 7. Comparison of the fitting models for the fluorescence lifetime analysis of shortT560, midT560 and longT560 in different cell lines.** Means of fluorescence lifetime T of shortT560, midT560 and longT560 extracted from fitting with single-exponential model, as well as of the two fluorescence lifetime components $T_1$ and $T_2$ and intensity-weighted average fluorescence lifetime $T_i$ from bi-exponential model in (**a**) HEK293T cells, (**b**) HeLa cells and (**c**) U2OS cells. N cells from two biological replicates were analyzed (the number of cells N is indicated on the right). The black lines represent the means of the two biological replicates. Each dot corresponds to the mean of each biological replicate (the lifetimes of individual cells are not shown). On the right is shown the distribution of goodness of fit (reduced $\chi^2$ or chi square) of mono- and bi-exponential models. Each cell is color-coded according to the biological replicate it came from. The solid circles correspond to the mean of each biological replicate. The black line represents the mean ± SD of the two biological replicates.

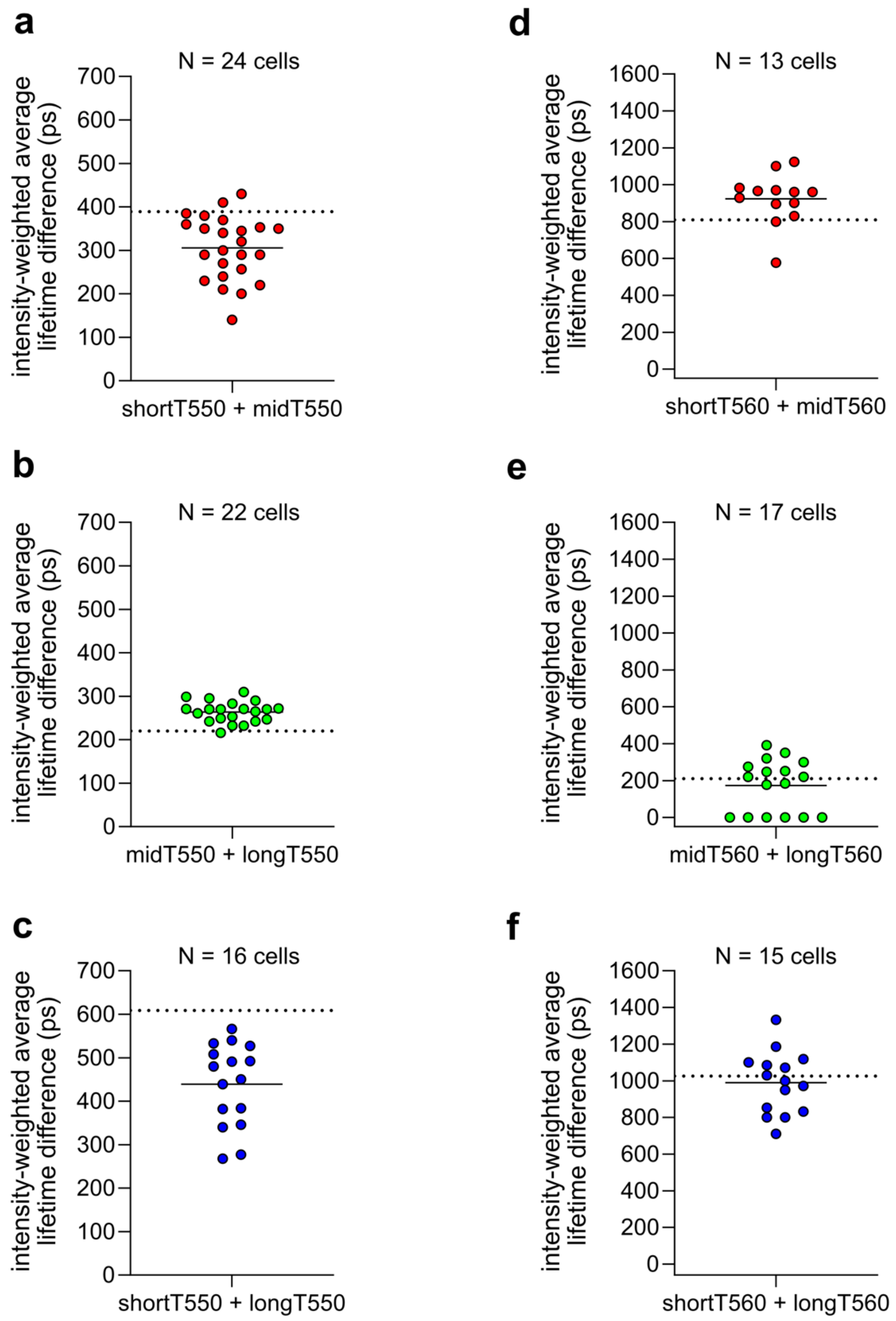


**Supplementary Figure 8. Pairwise intensity-weighted lifetime differences of short-FAST, midT-FAST and longT-FAST.** Each dot corresponds to the differences in intensity-weighted between mito-shortT550 and H2B-midT550 (**a**), mito-midT550 and H2B-longT550 (**b**), mito-shortT550 and H2B-longT550 (**c**), mito-short560 and H2B-midT560 (**d**), mito-midT560 and H2B-longT560 (**e**) and mito-shortT560 and H2B-longT560 (**f**). The dotted line indicates the intensity-weighted average lifetime difference between H2B-FAST variants determined in HeLa cells during the individual variants characterization (see also **Fig. 1**). The number of cells N (from at least three biological replicates) is indicated for each set of experiments.

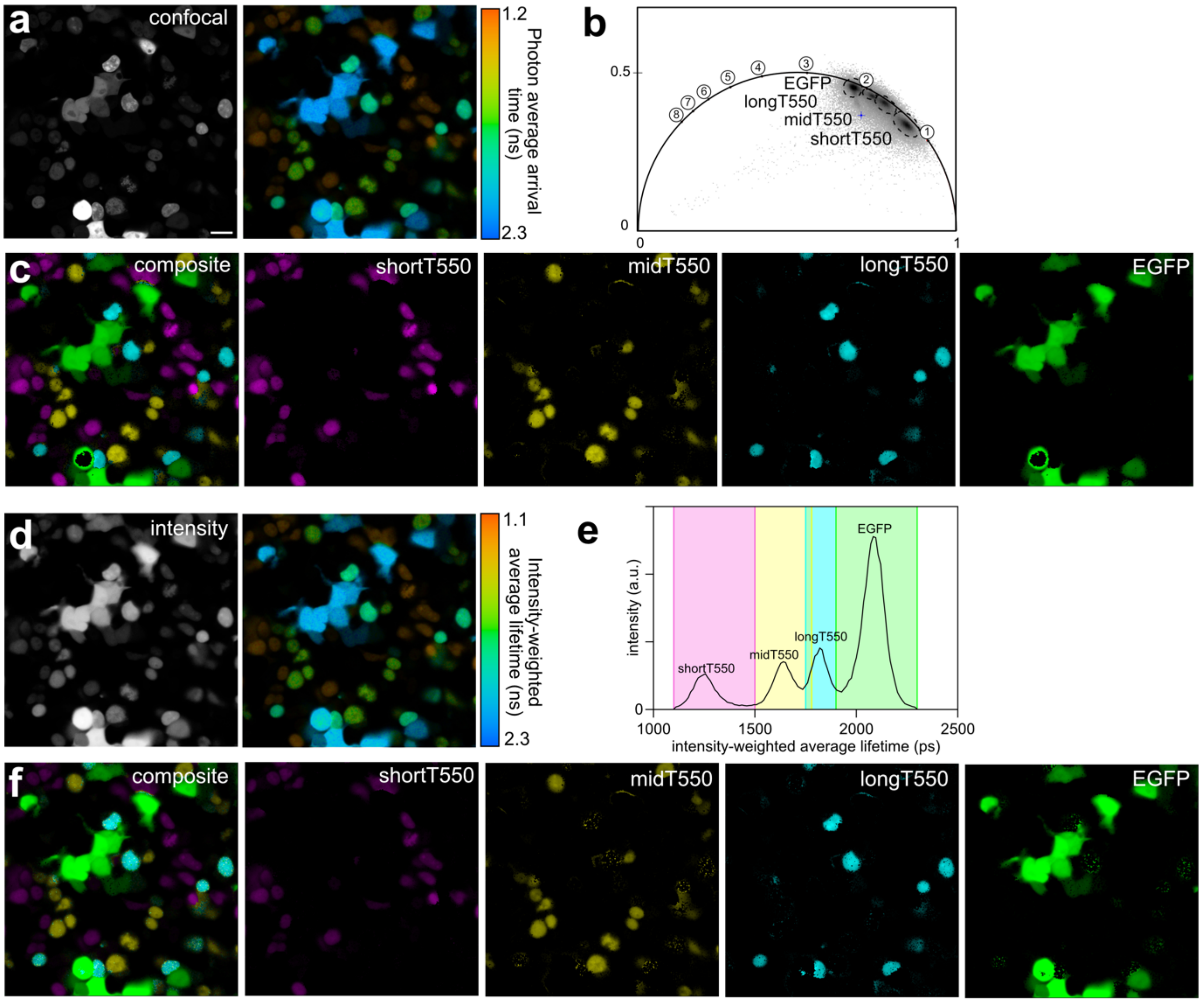


**Supplementary Figure 9. Fluorescence lifetime multiplexing in the green channel with shortT550, midT550, longT550 and EGFP.** (**a**) Confocal micrographs and equivalent average arrival time of photons color-coded image of a mix of HEK293T cells expressing H2B-shortT-FAST, H2B-midT-FAST, H2B-longT-FAST and cytosolic EGFP and labeled with 10 μM HBR-2,5DM. Scale bars, 20 μm. Excitation 488 nm / detection window 508-570 nm. (**b**) Corresponding phasor representation showing four separable clusters assigned to shortT550, midT550, longT550 and EGFP (time in ns is shown on the universal circle). (**c**) Phasor based separation of shortT550, midT550, longT550 and EGFP. Are shown the four individual isolated populations and a composite of all. (**d**) Intensity image micrographs and equivalent intensity-weighted average lifetime (bi-exponential fit) of a mix of HEK293T cells expressing H2B-shortT-FAST, H2B-midT-FAST, H2B-longT-FAST and cytosolic EGFP and labeled with 10 μM HBR-2,5DM. (**e**) Corresponding intensity-weighted average lifetime histogram showing four separated peaks, assigned to shortT550, midT550, longT550 and EGFP. (**f**) Intensity-weighted based separation of shortT550, midT550, longT550 and EGFP. Are shown the four individual isolated populations and a composite of all. Representative results of six fields of view from two biological replicates.

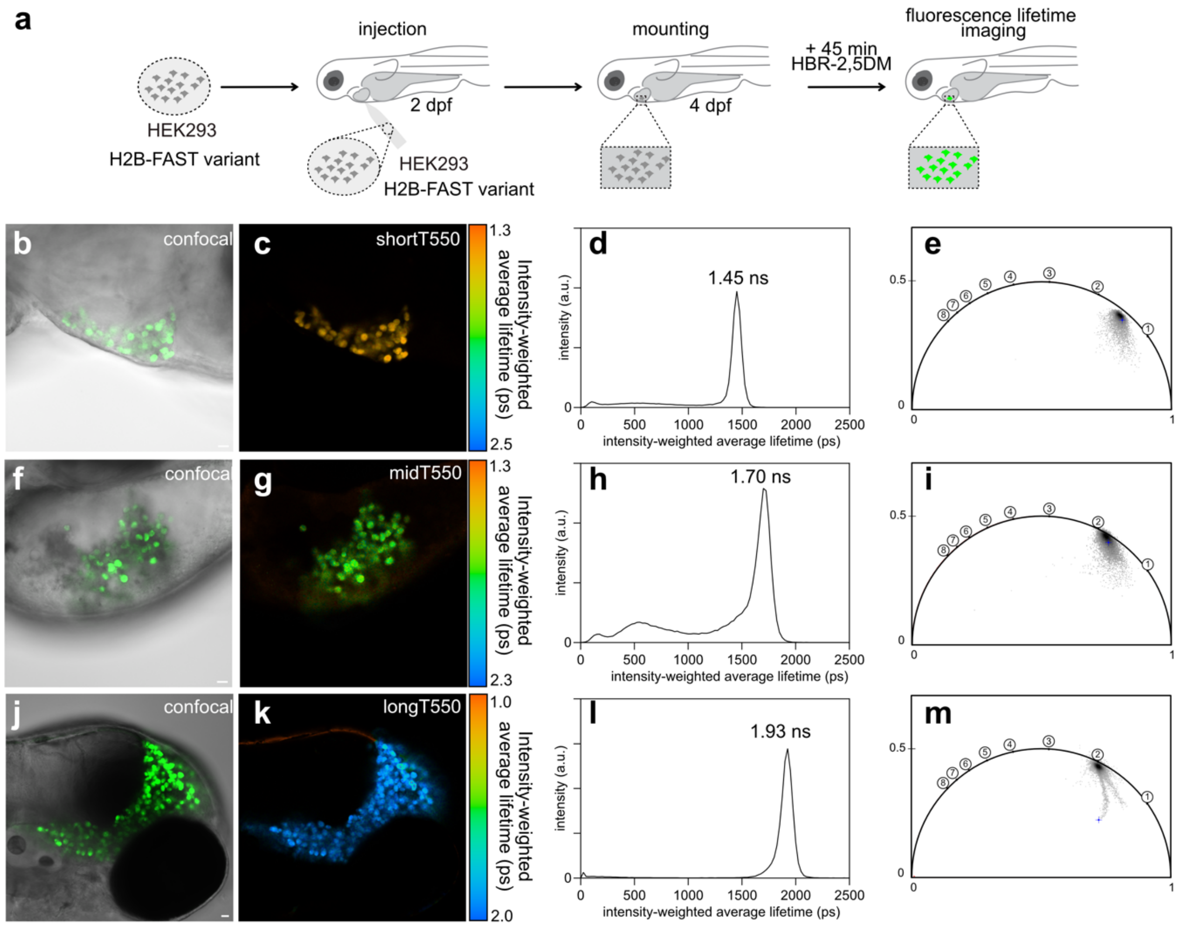


**Supplementary Figure 10. Fluorescence lifetime characterization of shortT550, midT550 and longT550 in zebrafish larvae. (a)** Mammalian HEK293T were transfected with plasmid encoding either (**b-e**) H2B-shortT-FAST, (**f-i**) H2B-midT-FAST and (**j-m**) H2B-longT-FAST. After 24h, cell populations were injected either in the vitellus or in the developing brain of 2 dpf zebrafish larvae. Larvae were imaged at 4 dpf after 45 min incubation with HBR-2,5DM. (**b,f,j**) Confocal micrographs. Scale bars, 20 µm. Excitation 488 nm / detection window 508-570 nm. (**c,g,k**) Intensity-weighted average lifetime color coded images. (**d,h,l**) Intensity-weighted lifetime histogram obtained by fitting the fluorescence decay data with a bi-exponential model. (**e,I,m**) Phasor plots (time in ns is shown on the universal circle). Experiments were repeated three times with similar results.

**Supplementary Table 1. FAST variants properties with HBR-2,5DM**

| | $K_D$ (μM) | $\lambda_{abs}$ (nm) | ε ($mM^{-1}.cm^{-1}$) | $\lambda_{em}$ (nm) | FQY (%) | $T_i$ (ns) | $k_r$ ($\times 10^8$ $s^{-1}$) | $k_{nr}$ ($\times 10^8$ $s^{-1}$) |
|---|---|---|---|---|---|---|---|---|
| pFAST | 0.01 | 498 | 52 | 549 | 0.30 | 1.79 | 1.68 | 3.91 |
| greenFAST | 0.12 | 494 | 54 | 551 | 0.25 | 1.34 | 1.87 | 5.60 |
| oFAST | 0.01 | 496 | 51 | 547 | 0.29 | 1.72 | 1.69 | 4.13 |
| TsiA-FAST | 0.01 | 498 | 52 | 549 | 0.34 | 1.91 | 1.78 | 3.46 |

Abbreviations are as follows: $K_D$ thermodynamic dissociation constant, $\lambda_{abs}$ wavelength of maximal absorption, ε molar absorptivity at $\lambda_{abs}$ (standard error is typically 10%), $\lambda_{em}$ wavelength of maximal emission, FQY fluorescence quantum yield. $T_i$ intensity-weighted average fluorescence lifetime (in HEK293T cells), $k_r$ radiative rate constant, $k_{nr}$ non radiative rate constant. The constants $k_r$ and $k_{nr}$ were computed using the FQY and $T_i$ values.

**Supplementary Table 2. FAST variants properties with HBR-3,5DM**

| | $K_D$ (μM) | $\lambda_{abs}$ (nm) | ε ($mM^{-1}.cm^{-1}$) | $\lambda_{em}$ (nm) | FQY (%) | $T_i$ (ns) | $k_r$ ($\times 10^8$ $s^{-1}$) | $k_{nr}$ ($\times 10^8$ $s^{-1}$) |
|---|---|---|---|---|---|---|---|---|
| pFAST | 0.01 | 502 | 49 | 561 | 0.44 | 2.94 | 1.50 | 1.90 |
| greenFAST | 0.83 | 498 | 50 | 559 | 0.38 | 2.12 | 1.79 | 2.92 |
| oFAST | 0.014 | 502 | 48 | 561 | 0.43 | 2.98 | 1.44 | 1.91 |
| TsiA-FAST | 0.012 | 500 | 45 | 561 | 0.47 | 3.05 | 1.54 | 1.74 |

Abbreviations are as follows: $K_D$ thermodynamic dissociation constant, $\lambda_{abs}$ wavelength of maximal absorption, ε molar absorptivity at $\lambda_{abs}$ (standard error is typically 10%), $\lambda_{em}$ wavelength of maximal emission, FQY fluorescence quantum yield. $T_i$ intensity-weighted average fluorescence lifetime (in HEK293T cells), $k_r$ radiative rate constant, $k_{nr}$ non radiative rate constant. The constants $k_r$ and $k_{nr}$ were computed using the FQY and $T_i$ values.

| Vector | ORF | ORF sequences |
| --- | --- | --- |
| pAG109 | H2B-<br>FAST-<br>cMyc | atgcccgaacctgcgaagtcagcgcccgctcccaaaaaggctctaaaaaagctgtc<br>gccaagacccagaagaagggggataagaaaaggcgtaagaccaggaaagagagt<br>tacgccatttacgtgtacaaagtactaaaacaagtccacccggacactggcatctcctca<br>aaggcgatgggcattatgaactcatttgtaaacgacatcttcgagcgcatcgccggaga<br>agcgtcgcgcctggcgcattacaacaagcgctccactatcacatcccgggagatccag<br>acggccgtgcgcctgctcctgcccggagaactggccaaacacgctgtgtctgagggca<br>caaaggccgtgaccaagtacaccagctccaagggcggaggctccggaggcggatct<br>gccaccatggagcatgttgcctttggcagtgaggacatcgagaacactctggccaaaat<br>ggacgacggacaactggatgggttggcctttggcgcaattcagctcgatggtgacggg<br>aatatcctgcagtacaatgctgctgaaggagacatcacaggcagagatcccaaacag<br>gtgattgggaagaacttcttcaaggatgttgcacctggaacggattctcccgagttttacg<br>gcaaattcaaggaaggcgtagcgtcagggaatctgaacaccatgttcgaatggatgat<br>accgacaagcaggggaccaaccaaggtcaaggtgcacatgaagaaagccctttccg<br>gtgacagctattgggtctttgtgaaacgggtgggatccgaacaaaagcttatttctgaaga<br>ggacttg |
| pAG374 | H2B-<br>greenFAS<br>T-cMyc | atgcccgaacctgcgaagtcagcgcccgctcccaaaaaggctctaaaaaagctgtc<br>gccaagacccagaagaagggggataagaaaaggcgtaagaccaggaaagagagt<br>tacgccatttacgtgtacaaagtactaaaacaagtccacccggacactggcatctcctca<br>aaggcgatgggcattatgaactcatttgtaaacgacatcttcgagcgcatcgccggaga<br>agcgtcgcgcctggcgcattacaacaagcgctccactatcacatcccgggagatccag<br>acggccgtgcgcctgctcctgcccggagaactggccaaacacgctgtgtctgagggca<br>caaaggccgtgaccaagtacaccagctccaagggcggaggctccggaggcggatct<br>gccaccatggagcatgttgcctttggcagtgaggacatcgagaacactctggccaaaat<br>ggacgacgaacaactggatgggttggcctttggcgcaattcagctcgatggtgacggg<br>aatatcctgcagtacaatgctgctgaaggagacatcacaggcagagatcccaaacag<br>gtgattgggaagaacttcttcaaggatgttgcaactggaacggattctcccgagttttacc<br>gcaaattcaaggaaggcgtagcgtcagggaatctgaacaccatgttcgaatggatgat<br>accgacaagcaggggaccaaccaaggtcaaggtgcacatgaagaaagccctttccg<br>gtgacagctattgggtctttgtgaaacgggtgggatccgaacaaaagcttatttctgaaga<br>ggacttg |
| pAG472 | H2B-<br>iFAST-<br>cMyc | atgcccgaacctgcgaagtcagcgcccgctcccaaaaaggctctaaaaaagctgtc<br>gccaagacccagaagaagggggataagaaaaggcgtaagaccaggaaagagagt<br>tacgccatttacgtgtacaaagtactaaaacaagtccacccggacactggcatctcctca<br>aaggcgatgggcattatgaactcatttgtaaacgacatcttcgagcgcatcgccggaga<br>agcgtcgcgcctggcgcattacaacaagcgctccactatcacatcccgggagatccag<br>acggccgtgcgcctgctcctgcccggagaactggccaaacacgctgtgtctgagggca<br>caaaggccgtgaccaagtacaccagctccaagggcggaggctccggaggcggatct<br>gccaccatggagcatgttgcctttggcagtgaggacatcgagaacactctggccaaaat<br>ggacgacggacaactggatgggttggcctttggcgcaattcagctcgatggtgacggg<br>aatatcctgcagtacaatgctgctgaaggagacatcacaggcagagatcccaaacag<br>gtgattgggaagaacttcttcaaggatgttgcacctggaacggattctcccgagttttacg<br>gcaaattcaaggaaggcgtagcgtcagggaatctgaacaccatgttcgaatggatgat<br>accgacaagcaggggaccaaccaaggtcaagatacacatgaagaaagccctttccg<br>gtgacagctattgggtctttgtgaaacgggtgggatccgaacaaaagcttatttctgaaga<br>ggacttg |

| | | |
|---|---|---|
| pAG657 | H2B-pFAST-cMyc | atgcccgaacctgcgaagtcagcgcccgctcccaaaaaaggctctaaaaaagctgtcgccaagacccagaagaaggggataagaaaaggcgtaagaccaggaaagagagttacgccatttacgtgtacaaagtactaaaacaagtccacccggacactggcatctcctcaaaggcgatgggcattatgaactcatttgtaaacgacatcttcgagcgcatcgccggagaagcgtcgcgcctggcgcattacaacaagcgctccactatcacatcccgggagatccagacggccgtgcgcctgctcctgcccggagaactggccaaacacgctgtgtctgagggcacaaaggccgtgaccaagtacaccagctccaagggcggaggctccggaggcggatctgccaccatggagcatgttgcctttggcagtgaggacatcgagaacactctggccaatatggacgacgaacaactggataggttggcctttggcgtaattcagctcgatggtgacggaatatcctgctgtacaatgctgctgaaggggacatcactggcagagatcccaaacaggtgattgggaagaacttcttcaaggatgttgcacctggaacggatactcccgagttttacggcaaattcaaggaaggcgcagcgtcagggaatctgaacaccatgttcgaatggacgataccgacaagcaggggaccaaccaaggtcaaggtgcacttgaagaaagccctttccggtgacagatattgggtctttgtgaaacgggtgggatccgaacaaaagcttatttctgaagaggacttg |
| pAG658 | H2B-tFAST-cMyc | atgcccgaacctgcgaagtcagcgcccgctcccaaaaaaggctctaaaaaagctgtcgccaagacccagaagaaggggataagaaaaggcgtaagaccaggaaagagagttacgccatttacgtgtacaaagtactaaaacaagtccacccggacactggcatctcctcaaaggcgatgggcattatgaactcatttgtaaacgacatcttcgagcgcatcgccggagaagcgtcgcgcctggcgcattacaacaagcgctccactatcacatcccgggagatccagacggccgtgcgcctgctcctgcccggagaactggccaaacacgctgtgtctgagggcacaaaggccgtgaccaagtacaccagctccaagggcggaggctccggaggcggatctgccaccatggagcatgttgcctttggcagtgaggacatcgagaacactctggccaaaatggacgacggacaactggataggttggcctttggcgcaattcagctcgatggtgacgggaatatcctgaagtacaatgctgctgaaggagacatcacaggcagagatcccaaacaggtgattgggaagaacttcttcaaggatgttgcacctggaacggatactcccgagttttacggcaaattcaaggaaggcgtatcgtcagggaatctgaacaccatgttcgaatgggcgataccgacaagcaggggaccaaccaaggtcaaggtgcatttgaagaaagccctttccggtgacagatattgggtctttgtgaaacgggtgggatccgaacaaaagcttatttctgaagaggacttg |
| pAG659 | H2B-oFAST-cMyc | atgcccgaacctgcgaagtcagcgcccgctcccaaaaaaggctctaaaaaagctgtcgccaagacccagaagaaggggataagaaaaggcgtaagaccaggaaagagagttacgccatttacgtgtacaaagtactaaaacaagtccacccggacactggcatctcctcaaaggcgatgggcattatgaactcatttgtaaacgacatcttcgagcgcatcgccggagaagcgtcgcgcctggcgcattacaacaagcgctccactatcacatcccgggagatccagacggccgtgcgcctgctcctgcccggagaactggccaaacacgctgtgtctgagggcacaaaggccgtgaccaagtacaccagctccaagggcggaggctccggaggcggatctgccaccatggagcatgttgcctttggcagtgaggacatcgagaacactctggccaaaatggacgacggacaactggatgggttggcctttggcgcaattcagctcgatggtgacgggaatatcctgctgtacaatgctgctgaaggagacatcacaggcagagatcccaaacaggtgattgggaagaacttcttcaaggatgttgcacctggaacgaattctcccgagttttacggcaaattcaaggaaggcatagcgtcagggaatctgaacaccatgttcgaatggatgataccgacaagcaggggaccaaccaaggtcaaggtgcacttgaagaaagccctttccggtgacagatattgggtctttgtgaaacgggtgggatccgaacaaaagcttatttctgaagaggacttg |
| pAG1448 | H2B-TsiA-FAST-cMyc | atgcccgaacctgcgaagtcagcgcccgctcccaaaaaaggctctaaaaaagctgtcgccaagacccagaagaaggggataagaaaaggcgtaagaccaggaaagagagttacgccatttacgtgtacaaagtactaaaacaagtccacccggacactggcatctcctcaaaggcgatgggcattatgaactcatttgtaaacgacatcttcgagcgcatcgccggagaagcgtcgcgcctggcgcattacaacaagcgctccactatcacatcccgggagatccagacggccgtgcgcctgctcctgcccggagaactggccaaacacgctgtgtctgagggcacaaaggccgtgaccaagtacaccagctccaagggcggaggctccggaggcggatctgccaccatggagctgctgagcttcggcgccgacaacatcgagaacagcctggccaagatgagcaagggcgacctgaacaagctggccttcggcgccatccagctgaacgcccagggcaagatcctgcagtacaacgccgccgagggcgacatcaccggcagaaagccca |

| | | |
|---|---|---|
| | | ccgaggtgatcggcaagaacttcttcctggaggtggcccccggcaccaacagaaccg<br>agttcaagggcagattcgaccagggcatcaagagcggcaacctgaacaccatgttcg<br>agtggatgatccccaccagcagaggccccaccaaggtgaaggtgcacatgaagaag<br>gccctggtggacgacacctactgggtgttcgtgaagagagtgggatccgaacaaaagc<br>ttatttctgaagaggacttg |
| pAG372 | Mito-greenFAST-cMyc | atgtccgtcctgacgccgctgctgctgcggggcttgacaggctcggcccggcggctccc<br>agtgccgcgcgccaagatccattcgttgagatctgccaccatggagcatgttgcctttggc<br>agtgaggacatcgagaacactctggccaaaatggacgacgaacaactggatgggttg<br>gcctttggcgcaattcagctcgatggtgacgggaatatcctgcagtacaatgctgctgaa<br>ggagacatcacaggcagagatcccaaacaggtgattgggaagaacttcttcaaggat<br>gttgcaactggaacggattctcccgagttttaccgcaaattcaaggaaggcgtagcgtca<br>gggaatctgaacaccatgttcgaatggatgataccgacaagcaggggaccaaccaag<br>gtcaaggtgcacatgaagaaagccctttccggtgacagctattgggtctttgtgaaacgg<br>gtgggatccgaacaaaagcttatttctgaagaggacttg |
| pAG673 | Mito-oFAST-cMyc | atgtccgtcctgacgccgctgctgctgcggggcttgacaggctcggcccggcggctccc<br>agtgccgcgcgccaagatccattcgttgagatctgccaccatggagcatgttgcctttggc<br>agtgaggacatcgagaacactctggccaaaatggacgacggacaactggatgggttg<br>gcctttggcgcaattcagctcgatggtgacgggaatatcctgctgtacaatgctgctgaag<br>gagacatcacaggcagagatcccaaacaggtgattgggaagaacttcttcaaggatgt<br>tgcacctggaacgaattctcccgagttttacggcaaattcaaggaaggcatagcgtcag<br>ggaatctgaacaccatgttcgaatggatgataccgacaagcaggggaccaaccaagg<br>tcaaggtgcacttgaagaaagccctttccggtgacagatattgggtctttgtgaaacgggt<br>gggatccgaacaaaagcttatttctgaagaggacttg |
| | Hbol-FAST | atggagaccgtgagattcggcggcgacgacatcgagaacagcctggccaagatgga<br>cgacaagaagctggacgagctggccttcggcgccatccagctggacgccaacggca<br>agatcatccagtacaacgccgccgagggcggcatcaccggcagagaccccaagag<br>cgtgatcggcaagaacttcttcaccgaggtggcccccggcacccagagcaaggagttc<br>cagggcagattcaaggagggcgtgagcagcggcgagctgaacaccatgttcgagtg<br>gatgatccccaccagcagaggccccaccaaggtgaaggtgcacatgaagaaggcc<br>atcagcggcgacacctactggatcttcgtgaagagactg |
| | HspG-FAST | atggagaccgtgagattcggcggcgacgacatcgagaacgccctggccaacatgga<br>cgacaagaagctggacaccctggccttcggcgccatccagctggacgccaacggca<br>agatcatccagtacaacgccgccgagggcggcatcaccggcagagaccccaagag<br>cgtgatcggcaagaacttcttcaccgacgtggcccccggcacccagagcaaggagttc<br>cagggcagattcaaggagggcgtgaagaacggcgacctgaacaccatgttcgagtg<br>gatgatccccaccagcagaggccccaccaaggtgaaggtgcacatgaagaaggccc<br>tgagcggcgacaccttctggatcttcgtgaagagactg |
| | RspA-FAST | atggagaccgtgagattcggcggcgacgacatcgagaacagcctggccaagatgga<br>cgacaaggccctggacaagctggccttcggcgccatccagctggacggcaacggca<br>agatcatccactacaacgccgccgagggcaccatcaccggcagagaccccaagacc<br>gtgatcggcaagaacttcttcaccgacgtggcccccggcacccagagcaaggagttcc<br>agggcagattcaaggagggcgtgcagaagggcgacctgaacaccatgttcgagtgg<br>atgatccccaccagcagaggccccaccaaggtgaaggtgcacatgaagaaggccat<br>gacggcgacagcttctggatcttcgtgaagagactg |
| | Ilo-FAST | atggagatcgtgcagttcggcagcgacgacatcgagaacaccctgagcaagatgagc<br>gacgacaagctgaacgacatcgccttcggcgccatccagctggacgccagcggcaa<br>gatcatccagtacaacgccgccgagggcgacatcaccggcagagaccccggcgcc<br>gtggtgggcaagaacttcttcaacgaggtggcccccggcaccaacagccccgagttca<br>agggcagattcgacgagggcgtgaagaacggcaacctgaacaccatgttcgagtgg<br>atgatccccaccagcagaggccccaccaaggtgaaggtgcacatgaagaaggccct<br>gaccggcgacacctactgggtgttcgtgaagagactg |
| | RsA-FAST | atggagatgatcaagttcggccaggacgacatcgagaacgccatggccgacatgggc<br>gacgcccagatcgacgacctggccttcggcgccatccagctggacgagaccggcacc<br>atcctggcctacaacgccgccgagggcgagctgaccggcagaagcccccaggacgt<br>gatcggcaagaacttcttcaaggacatcgcccccggcaccgacaccgaggagttcgg |

| | | |
|---|---|---|
| | | cggcagattcagagagggcgtggccaacggcgacctgaacgccatgttcgagtggat<br>gatccccaccagcagaggccccaccaaggtgaaggtgcacatgaagagagccatca<br>ccggcgacagctactggatcttcgtgaagagagtg |
| pAG1367 | H2B-<br>emIRFP67<br>0-cMyc | atgcccgaacctgcgaagtcagcgcccgctcccaaaaaggctctaaaaagctgtc<br>gccaagacccagaagaagggggataagaaaaggcgtaagaccaggaaagagagt<br>tacgccatttacgtgtacaaagtactaaaacaagtccacccggacactggcatctcctca<br>aaggcgatgggcattatgaactcatttgtaaacgacatcttcgagcgcatcgccggaga<br>agcgtcgcgcctggcgcattacaacaagcgctccactatcacatcccgggagatccag<br>acggccgtgcgcctgctcctgcccggagaactggccaaacacgctgtgtctgagggca<br>caaaggccgtgaccaagtacaccagctccaagggcggaggctccggaggcggatct<br>gccaccatggcggaaggatccgtcgccaggcagcctgacctcttgacctgcgaacatg<br>aagagatccacctcgccggctcgatccagccgcatggcgcgcttctggtcgtcagcga<br>acatgatcatcgcgtcatccaggccagcgccaacgccgcggaatttctgaatctcggaa<br>gcgtactcggcgttccgctcgccgagatcgacggcgatctgttgatcaagatcctgccgc<br>atctcgatcccaccgccgaaggcatgccggtcgcggtgcgctgccggatcggcaatcc<br>ctctacggagtactgcggtctgatgcatcggcctccggaaggcgggctgatcatcgaac<br>tcgaacgtgccggcccgtcgatcgatctgtcaggcacgctggcgccggcgctggagcg<br>gatccgcacggcgggttcactgcgcgcgctgtgcgatgacaccgtgctgctgtttcagca<br>gtgcaccggctacgaccgggtgatggtgtatcgtttcgatgagcaaggccacggcctgg<br>tattctccgagtgccatgtgcctgggctcgaatcctatttcggcaaccgctatccgtcgtcg<br>actgtcccgcagatggcgcggcagctgtacgtgcggcagcgcgtccgcgtgctggtcg<br>acgtcacctatcagccggtgccgctggagccgcggctgtcgccgctgaccgggcgcg<br>atctcgacatgtcgggctgcttcctgcgctcgatgtcgccgtgccatctgcagttcctgaag<br>gacatgggcgtgcgcgccaccctggcggtgtcgctggtggtcggcggcaagctgtggg<br>gcctggttgtctgtcaccattatctgccgcgcttcatccgtttcgagctgcgggcgatctgca<br>aacggctcgccgaaaggatcgcgacgcggatcaccgcgcttgagagcggatccgaa<br>caaaagcttatttctgaagaggacttg |
